# A semi-supervised Bayesian mixture modelling approach for joint batch correction and classification

**DOI:** 10.1101/2022.01.14.476352

**Authors:** Stephen Coleman, Kath Nicholls, Xaquin Castro Dopico, Gunilla B. Karlsson Hedestam, Paul D.W. Kirk, Chris Wallace

**Affiliations:** MRC Biostatistics Unit, University of Cambridge, U.K.; Cambridge Institute of Therapeutic Immunology & Infectious Disease, University of Cambridge, U.K.; Department of Microbiology, Tumor and Cell Biology, Karolinska Institutet, Sweden; Cancer Research U.K. Cambridge Centre, Ovarian Cancer Programme, University of Cambridge, U.K.

**Keywords:** SARS-CoV-2, ELISA, Mixture model, Batch correction, Bayes, Assay data, Classification

## Abstract

Systematic differences between batches of samples present significant challenges when analysing biological data. Such *batch effects* are well-studied and are liable to occur in any setting where multiple batches are assayed. Many existing methods for accounting for these have focused on high-dimensional data such as RNA-seq and have assumptions that reflect this. Here we focus on batch-correction in low-dimensional classification problems. We propose a semi-supervised Bayesian generative classifier based on mixture models that jointly predicts class labels and models batch effects. Our model allows observations to be probabilistically assigned to classes in a way that incorporates uncertainty arising from batch effects. By simultaneously inferring the classification and the batch-correction our method is more robust to dependence between batch and class than pre-processing steps such as ComBat. We explore two choices for the within-class densities: the multivariate normal and the multivariate *t*. A simulation study demonstrates that our method performs well compared to popular off-the-shelf machine learning methods and is also quick; performing 15,000 iterations on a dataset of 750 samples with 2 measurements each in 11.7 seconds for the MVN mixture model and 14.7 seconds for the MVT mixture model. We further validate our model on gene expression data where cell type (class) is known and simulate batch effects. We apply our model to two datasets generated using the enzyme-linked immunosorbent assay (ELISA), a spectrophotometric assay often used to screen for antibodies. The examples we consider were collected in 2020 and measure seropositivity for SARS-CoV-2. We use our model to estimate seroprevalence in the populations studied. We implement the models in C++ using a Metropolis-within-Gibbs algorithm, available in the R package `batchmix`. Scripts to recreate our analysis are at https://github.com/stcolema/BatchClassifierPaper.

## 1 Background

Many biological assays are performed across sets of samples or *batches*. When the number of samples exceeds the batch size, then it is common to notice *batch effects*, systematic differences between assay readouts from different batches which may affect both their mean and scale. This is a prevalent problem, that may be addressed in a variety of ways depending on the planned downstream analysis. In discussing available options for batch correction, we will use the term “batch effect” to mean differences between samples arising from between-batch technical factors in the experiment, and the term “class effect” to refer to biological differences arising due to samples coming from distinct biological classes. We consider settings in which the objective is to classify unlabelled samples into predefined classes.

To analyse class effects we should also account for the batch effects. One common approach is to first correct for batch effects as part of a pre-processing or data cleaning step (which might be as simple as zero-centring the data; i.e., transforming each batch to have a common mean), and then to apply standard classification models to the resulting “cleaned” data (e.g., 2, 27, 36). However, such two-step approaches have been found to increase false positive rates because they may induce correlation between the cleaned observations which is typically not accounted for in downstream analysis (25). Further, when batch is confounded with class effects (due to unbalanced representation of classes across batches) then naive adjustment which ignores known biological classes in the data can lead to incorrect conclusions (23), and methods for adjustment which preserve differences attributable to known classes can lead to false positive results (32). An alternative approach is to incorporate batch information directly into downstream analyses, for example as a covariate in regression-based approaches. It has been shown that mixed effects models which share information between batches produce better calibrated quantitative data than independent analyses of each batch (39). However, only a subset of analytical approaches have been adapted to accommodate batch effects (e.g., 31, 33, 22), and there has been a strong focus on high-dimensional settings (e.g., 18, 6, 1). Thus a need exists for a wider range of methods that can account for batch effects directly in low-dimensional data analysis.

Here we focus on the problem of assigning class labels using low-dimensional assay data generated across several batches. This is a common design in many assays that measure a small number of specific biomarkers such as enzyme-linked immunosorbent assay (ELISA) and flow cytometry data. If there are known classes in the population, then class-specific controls can be included in the assay, resulting in training examples for which the class labels are known. We are motivated in part by the specific problem of estimating seroprevalance of SARS-CoV-2 by classifying individuals into seropositive and seronegative classes at different points in time during the pandemic. Since batches tend to comprise samples collected at the same time point, and since seroprevalence is expected to vary through the course of the pandemic, we expect class membership to be imbalanced across batches – motivating the development of a joint classification and batch-correction model, rather than a 2-step approach. Insofar as we are aware, there is no appropriate method for classification using data with all of these characteristics.

To address this, we propose a semi-supervised Bayesian mixture model that explicitly models batch parameters and predicts class membership. Our method is semi-supervised as our model parameters are inferred using both the labelled and unlabelled data. In an iteration of our MCMC algorithm the unobserved class labels are inferred, then a subset of the data, *X_k_*, are associated with the *k^th^* class, and *X_k_* can be divided into unlabelled (u) and labelled (l) data, e.g., *X_k_* = [*X*_(*l*)*k*_, *X*_(*u*)*k*_], where the labels of *X*_(*l*)*k*_ are observed and the labels of *X*_(*u*)*k*_ are imputed. The class parameters are then inferred conditioning on the entirety of *X_k_*. This happens in each iteration of our sampler and the algorithm is run for at least as many iterations as it takes to converge. Semi-supervised Bayesian mixture models have had some success in biomedical applications, e.g. Crook et al. (10, 11). The Bayesian framework also allows our model to propagate the uncertainty arising from the batch effects to the class allocation probabilities for each item in the dataset. This provides a more complete quantification of the uncertainty in the final predictions, thereby enabling more informed interpretation.

This manuscript is organised as follows: in section 2 we describe our model; in section 3 we evaluate our model using simulated data, and compare to off-the-shelf machine learning methods; in section 4 we compare the same set of models on a gene expression dataset where the true class is known and we simulate batch-effects under two scenarios; and in section 5 we apply the proposed method to two ELISA studies of seroprevalence of SARS-CoV-2 in Stockholm (7) (section 5.1) and Seattle (12) (section 5.3). We then conclude our manuscript in section 6 with a discussion of the contribution, limitations, and possible extensions to our model.

## 2 Model

### 2.1 Notation

We consider a study that collects *P* measurements for each of *N* individuals to form a dataset *X* = (*X*_1_, …, *X_N_*), where *X_n_* = [*X*_*n*,1_,, …, *X_n,P_*]^⊤^ for all *n* ∈ {1, …, *N*}. We assume that each individual has an associated observed batch label 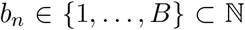, where *B* is the total number of batches, and we write *b* = [*b*_1_, …, *b_N_*]^⊤^ for the collection of all N batch labels. Note that as each individual belongs to a single batch, we assume that all *P* measurements for each individual are part of the same batch. We wish to predict class labels for each individual, and write *c* = [*c*_1_, …, *c_N_*]^⊤^ for the collection of all class labels. We assume that the number of classes, *K*, is known, so that each *c_n_* ∈ {1, …, *K*}. We assume that a subset of labels are observed and each class is represented in this subset.

### 2.2 Model specification

We use a *K*-component mixture model to describe the data *X*. The mixture model can be be written

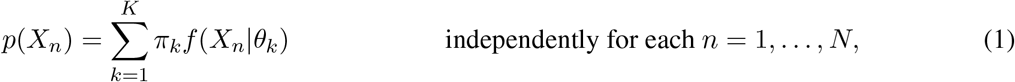

where π = [π_1_, …, π_*K*_]^⊤^ is the vector of component weights, *f*(·) is a parametric density function, and *θ_k_* are the parameters of the *k^th^* component. We assume each component describes a single and distinct class in the population and use the class labels to rewrite the model

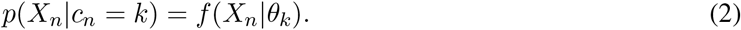

We then introduce batch-specific parameters, *z* = (*z*_1_, …, *z_B_*) and expand *f*(·) to accommodate these. Then conditioning on the observed batch label we have

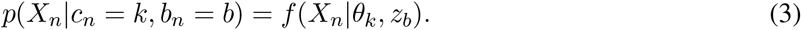

We focus on continuous data where each measurement has support across the entire real line. We consider the multivariate *t* density (**MVT**, density denoted *f_t_*(·)) and the multivariate normal (**MVN**, density denoted 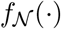) as choices for *f*, but depending on the situation other choices could be more relevant and our model is not inherently restricted to these. We use *z_b_* = (*m_b_, S_b_*), choosing *m_b_* to be a *P*-vector representing the shift in location due to the batch effects and *S_b_* to be a scaling matrix. We assume the observed location of *X_n_* is composed of a class-specific effect, *μ_k_*, and a batch-specific effect, *m_b_*, so (*X_n_*|*c_n_* = *k*, *b_n_* = *b*) = *μ_k_* + *m_b_* + *ϵ_n_*. Similarly we assume that the random noise, *ϵ_n_*, is subject to class and batch specific effects Σ_*k*_ and *S_b_* respectively.

More specifically, if we use a mixture of MVN densities, then our class parameters are *θ_k_* = (*μ_k_*, Σ_*k*_), where *μ_k_* is the *P*-dimensional mean vector and Σ_*k*_ is the *P* × *P* covariance matrix. We assume

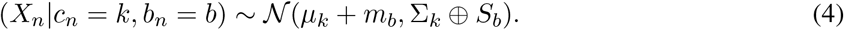

We define the operator ⊕ for a *P* × *P* matrix, *A*, and a diagonal matrix *B* of equal dimension, as:

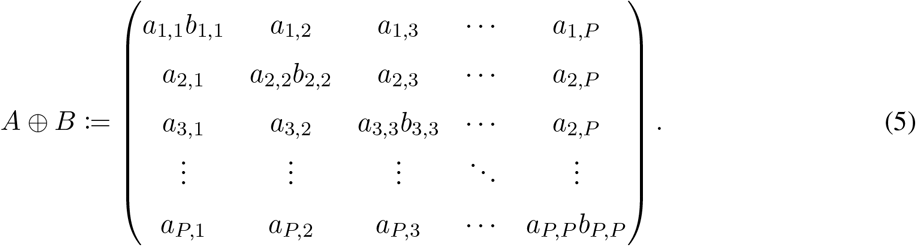

Similarly for a mixture of MVT densities, we assume

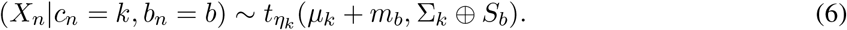

where *η_k_* is the class-specific degrees of freedom.

In the likelihood function, only the combinations of the class and batch parameters, *μ_k_* + *m_b_* and Σ_*k*_ ⊕ *S_b_*, are identifiable, and the values of the class and batch specific effects are not. However, we assume that we have some prior information about the relative orders of magnitude of the class and batch effects and encode this in an informative prior, reducing the problem of identifiability with this additional constraint. If the magnitude of the between-batch variability is similar to or greater than the true biological effect, then we suspect that any analysis of such a dataset is untenable, or at least that the data are not appropriate for our model.

The full hierarchical model can be found in section 1 of the supplementary material. Here we include the choice of prior distributions for the class and batch effects:

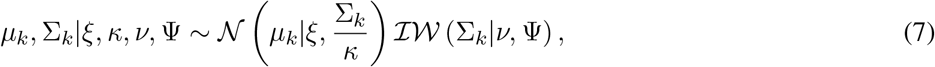

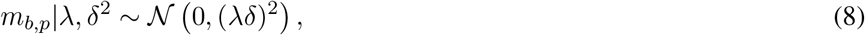

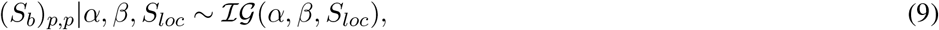

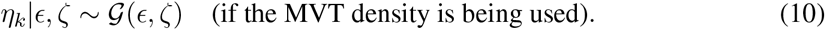

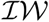 denotes the inverse-Wishart distribution, 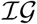 denotes the inverse-Gamma distribution with a shape *α*, rate *β* and location *S_loc_*, 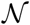 signifies the Gaussian distribution parameterised by a mean vector and a covariance matrix and 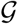 denotes the Gamma distribution parameterised by a shape and rate. An empirical Bayes approach is used to set the hyperparameters for the class mean and covariance (details are included in section 2 of the Supplementary material, these follow the suggestions of 14). The *δ*^2^ hyperparameter is set to the mean of the diagonal entries of the observed covariance in the data. *S_loc_* is set to 1.0 to ensure that the likelihood covariance matrix remains positive semi-definite. For the MVT mixture model, we choose the hyperparameters of the degrees of freedom to be *ϵ* = 20 *ζ* = 0.1 in line with suggestions from Juárez and Steel (20). This uninformative prior does not restrict *η_k_* to small values, and enables the MVT mixture model to approximate the MVN model if the data are truly Gaussian. The remaining hyperparameters (λ, *α* and *β*) are user-specified, and we explore the impact of different choices on the final inference in sections 5.1 and 5.3. We investigate the impact of 3 different values for each of these parameters, reflecting an informative or constrained prior, a flexible, uninformed prior, and a choice in the middle-ground.

Sampling the batch and class parameters allows us to derive a batch-corrected dataset, *Y*, in each iteration. We define the *p^th^* measurement for the *n^th^* sample in *Y* as

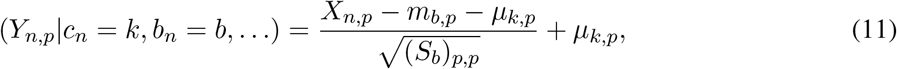

for all *n* = {1, …, *N*}, *p* = {1, …, *P*}. Note that *Y* will incorporate the uncertainty about the batch and class parameters, and the classification. This transformation is similar to the empirical Bayes batch correction suggested by Johnson et al. (19); however their method is a pre-processing step that is applied to each measurement in turn, whereas our model is jointly inferring class and batch effects and may be applied to the full dataset.

We perform inference using a Metropolis-within-Gibbs sampler as described in section 3 of the supplementary material.

## 3 Simulations

### 3.1 Simulation design

We wish to evaluate the performance of the MVN and MVT implementations of our model and compare these to the popular machine learning methods random forest (**RF**, 5), probabilistic support vector machine (**SVM**, 4) and logistic regression (without batch-correction, **LR**). We also compare our method to ComBat (19, 24), a popoular pre-processing batch-correction method. We apply ComBat to our data and then model the resulting dataset with the MVN and MVT mixture models and logistic regression. Finally, we consider the semi-supervised Bayesian mixture model with no batch-correction.

To achieve this, we generate 100 datasets in each of 9 different scenarios. In six of these, the data are generated from a mixture of MVN distributions and in one scenario the data is generated from a mixture of MVT distributions. In two scenarios, data are sampled from a a mixture of Poisson distributions. This count data is log-transformed and then Gaussian noise is added to ensure our model is strongly misspecified in the density choice. For each simulation we generate both a “batch-free” and an observed dataset. In seven of the scenarios, the data contains *P* = 2 measurements for each of *N* = 750 samples; in two scenarios we consider a higher feature space of *P* = 15. In all scenarios we consider *B* = 5 batches and *K* = 2 classes. The classes are not evenly represented, in seven scenarios the first class expected to contribute 75% of the samples with the remainder drawn from the second class, and class is independent of batch (and therefore a pre-processing step is expected to match our model). In two scenarios the class labels and batch of origin are dependent and pre-processing is expected to skew the inference. The batches are all expected to have equal numbers of samples except in the Varying batch size scenario. We randomly select which class labels are observed, sampling uniformly across the data indices, {1, …, *N*}. We expect one quarter of the labels to be observed, i.e. 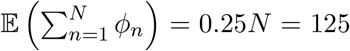. These labelled observations constitute the training set for the off-the-shelf methods.

More specifically, the nine simulation scenarios are:

- Base case: the generic, base scenario; all other scenarios are variations of this, using the same choices for all bar a subset of parameters, with this subset varied to define the specific scenario.
- Batch-free: similar to the Base case but no batch effects are present (i.e., *m_b_* = **0**_*P*_, *S_b_* = **I**).
- Varying batch effects: the Base case with more variance among the batch effects.
- Varying class representation: the classes are imbalanced across batches, i.e, the expected proportion of each class varies across batches (note that this is a slightly different generating model, the class weights are batch specific). The first two batches contain a larger proportion of samples from class 1, the third batch is balanced and the final two batches have a greater proportion of samples from class 2.
- Varying batch size: rather than equally sized batches, the batches have varying proportions of the total sample. The expected proportions are 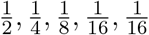.
- Multivariate t generated: the data are generated from a MVT mixture model rather than a MVN mixture model.
- Log-Poisson generated: the data are generated from a Poisson mixture model, log-transformed and then Gaussian noise is added.
- High-dimensional: *P* = 15 features are generated.
- Hardest: this combines the data-generating mechanism of the Log-Poisson generated scenario, the class and batch dependency of the varying class representation scenario and the dimensionality of the high-dimensional scenario.

A more detailed description of the generating models, along with visualisations of an example dataset for each scenario, are provided in section 4 of the supplementary material.

We use implementations of the machine learning methods available in `R` (34). For the RF this is the `randomForest` package (26), for the SVM we use the `kernlab` package (21), and for LR we use the base implementation of LR contained in the `glm` function. We use the default parameters in each method, bar the SVM where we set `prob.model = TRUE` to build a model for calculating class probabilities. The default for a classification SVM in this package uses a Gaussian Radial Basis kernel function.

We use the data with observed labels as the training set for each of these methods and those with unobserved labels as a test set. We record the time taken to train the model and to predict the outcome for the test set.

### 3.2 Results

We assessed within-chain convergence by calculating the Geweke statistic (16), and removed chains which failed the diagnostic test. We then selected chains with the highest median complete log-likelihood as representative for the simulation. We compared the models using the Brier score between allocation probability across classes and the true class (figure 1 A). We found that our mixture model performed better or at least as well as the ML methods across all two dimensional scenarios. When the data were generated from Gaussian distributions, the performance of the two versions of the mixture model performed very similarly. The MVT mixture model learned a large degree of freedom for each component, indicating that this behaves as an approximation of the Gaussian mixture model when appropriate (figure 2). In contrast, in the Multivariate *t* generated scenario, the performance of the MVN mixture model had greater variation in performance than in any other scenario. Figure 2 also shows that parameter estimates were consistent across chains. When there was no dependency between batch and class, the pre-processing batch-correction, ComBat, was as effective as the combined model (batchmix). However, when this dependency is introduced ComBat is problematic in keeping with results from Leek et al. (23), Li et al. (25), whereas batchmix is better protected from this confounding. bactchmix performed poorly in the high-dimensional scenarios, but this is potentially a problem with the sampler or the choice of priors as the model is no less well specified than in the Base case scenario. However, this shows that batchmix should not be casually applied in high-dimensional data.

**Figure 1:**
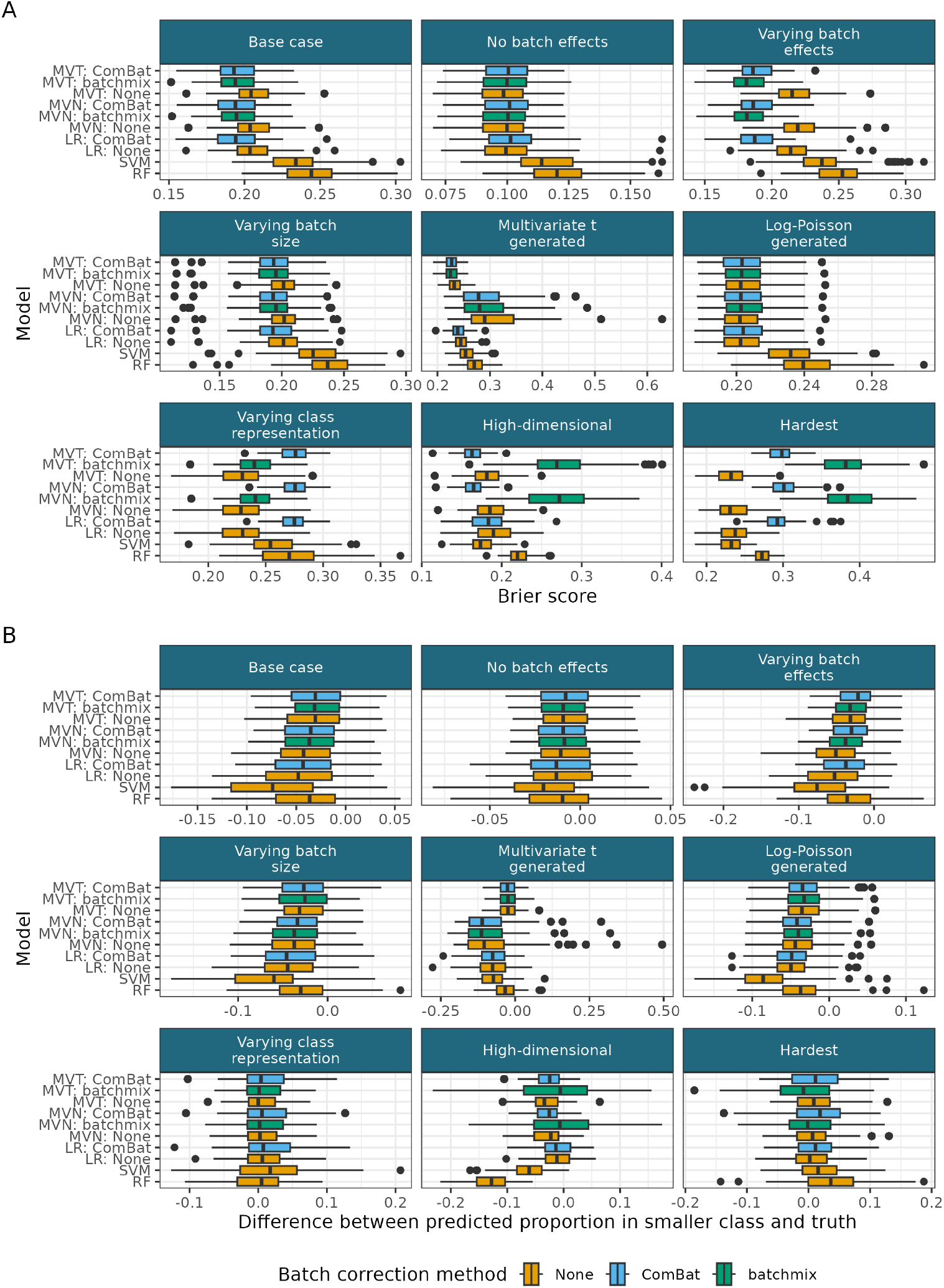
A) Brier score for the inferred allocation probability in the unlabelled data across simulations. B) The difference between the inferred proportion in class 2 and the true proportion of the data in class 2.

**Figure 2:**
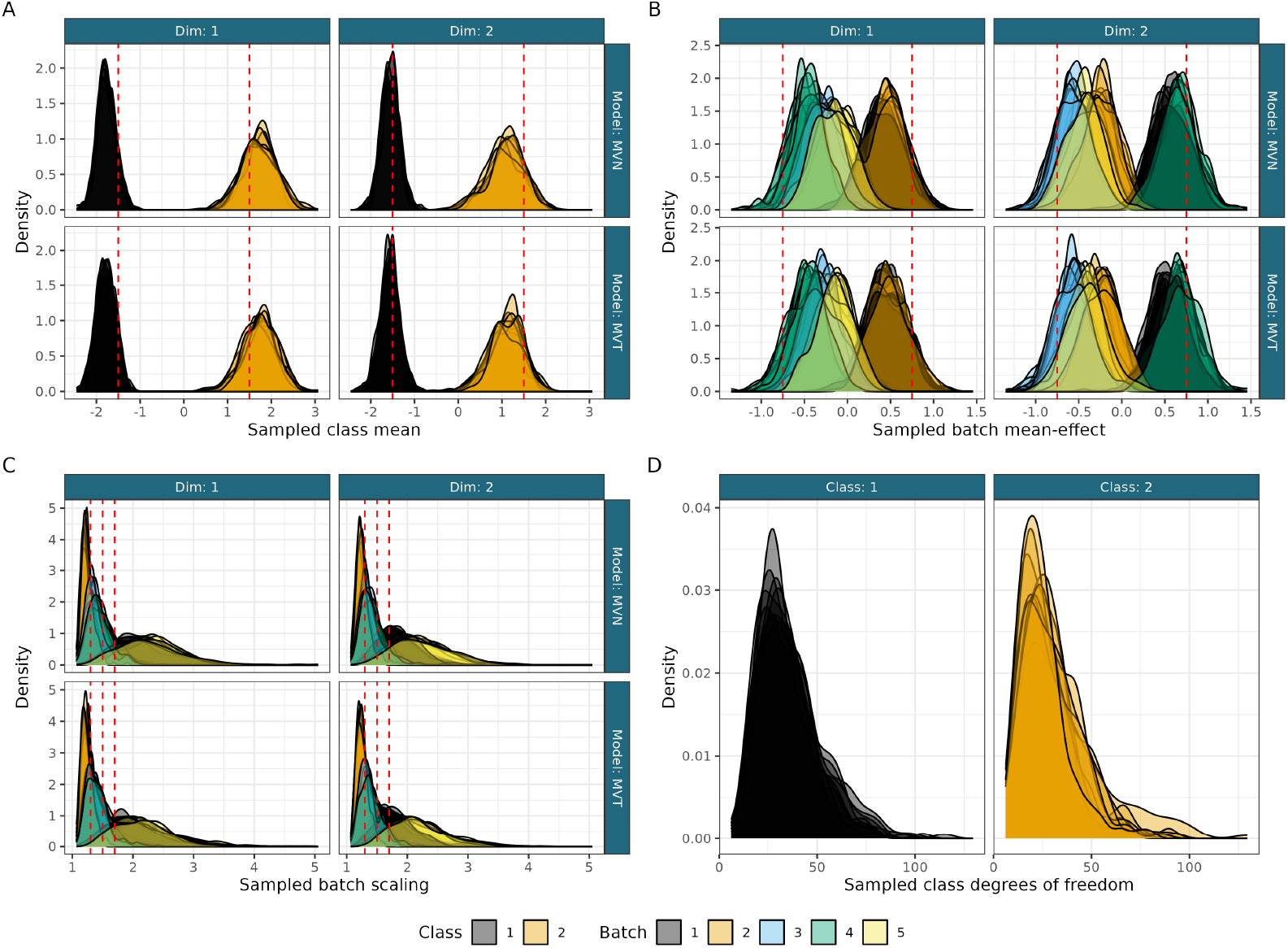
Sampled values for A) the class means, B) the batch mean-effect, C) the batch scaling effect and D) the class degrees of freedom for the well-behaved chains for the first simulated dataset in the Base case scenario. True values are shown by the dashed red vertical lines (as the data are generated from a MVN density there is no true degree of freedom, but larger values better approximate the MVN).

We also wanted a sense of how well our models would estimate the proportion of the classes in our simulations as an analogue to seroprevalence in our motivating example. We recorded the predicted proportion of the smaller class in the dataset, and compared the models’ estimate to the truth (figure 1 B). We found that the mixture models have a more narrow range in their estimates than the other models in the Base case, No batch effects, Varying batch effects and Varying batch size scenarios, with a similar range for the MVT mixture model in the other low-dimensaional scenarios indicating a more consistent behaviour than the other methods. The MVN mixture model exhibited good behaviour, except when misspecified as in the MVT generated data. We note that the mixture model’s median performance is either an under-estimate of the proportion of samples from the smaller class or to be centred on the true value. We also observed that when the batch effects were more varied and greater in magnitude, the SVM and RF had very long tails in their performance (Varying batch effects in figure 1 B).

The machine learning approaches and ComBat are all exceptionally fast, running in under a second. Running the MCMC for batchmix to converge took longer but was still reasonable. In the Base case scenario, 25,000 iterations had a median runtime of 11.7s for the MVN model and 14.7s for the MVT model. Full time results can be seen in figure 11 of the supplementary material.

## 4 Gene expression data

We consider a gene expression dataset for sorted peripheral blood cells (28) as an additional validation of the model. The dataset contains 84 CD14 cells, 24 CD16, 17 CD19, 24 CD4, and 25 CD8 cells. There are an additional unseparated 68 peripheral blood mononuclear cells (PBMC). We use the data as represented in the first four principal components to reduce the dimensionality of the data. We then scale the features by mean-centring them and transforming them to unit variance. We (artificially) introduce batch-effects under two scenarios a) the cell type and class are independent and b) the cell type and batch are correlated (figure 3 A and B; the generating models are described in section 5 of the supplementary material). We consider the same set of models as in the simulation study and aim to infer cell type for the data with hidden labels. We hide a random 80% of labels in each of ten folds with the restriction that each class has at least one known member. We compare model performance using the Brier score. The results are similar to the Base case scenario in the simulation study with the semi-supervised Bayesian mixture models performing the best. When there is no dependency between class and batch ComBat and batchmix are equivalent, but when this dependency is present ComBat suffers significantly. In this case doing no batch correction is the best strategy, followed closely by batchmix. In both scenarios the off-the-shelf supervised methods perform less well than the semi-supervised mixture models.

**Figure 3:**
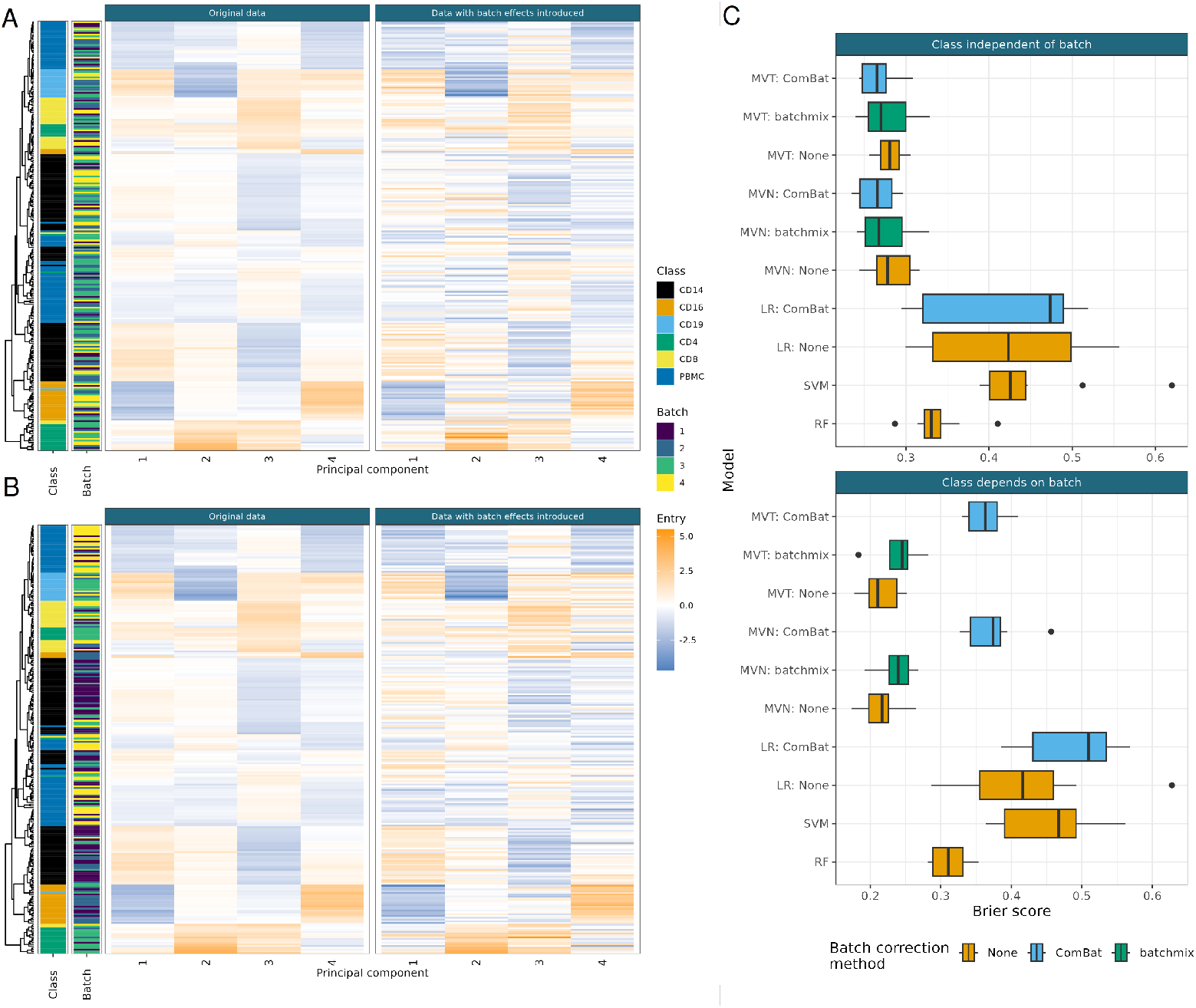
The data and model performance for the cell gene expression data. A) Facet on the left shows the original data, annotated by class and batch. The facet on the right shows the data after batch effects are introduced for class independent of batch. B) As A) but class is dependent on batch. C) Model performance across 10 folds under the Brier score for both scenarios.

## 5 ELISA data examples

ELISA is an immunological assay used to measure antibodies, antigens, proteins and glycoproteins, and normally involves a reaction that converts the substrate into a coloured product, the optical density (**OD**) which can be measured and is then used to determine the antigen concentration. One application is to assess seroprevalence of a disease within a population by measuring seropositivity of antibodies. It has a history of application to a wide range of diseases (e.g., 38, 3, 17, 30) and was used extensively to study seropositivity of antibodies to SARS-CoV-2 antigens used to estimate prevalence of cumulative infection and immunity (12, 29, 37). In such cases it is often possible to include known positive and negative controls as samples (these might be PCR-positive patients and historical samples collected before the pandemic began) and thus a subset of labels are observed.

We investigated the performance of our model on two recent examples of ELISA data, both from studies estimating seroprevalence of SARS-CoV-2. Based on the results from the simulations, we use the MVT as our choice of density, as it always matched or outperformed the MVN mixture in simulations (figure 1).

In the ELISA datasets we do not know the true seropositive status for the non-control data and cannot evaluate the model accuracy. Rather, we present these to demonstrate application of our model and highlight how diagnostic plots and results may be interpreted. In each case we run multiple chains and then use the sampled log-likelihood to assess within and across chain convergence.

Traditional analysis of ELISA data in seroprevalence studies makes dichotomous calls according to thresholds based on the sum of the sample mean and some number of standard deviations of the negative controls in each measurement. However, various choices of the number of standard deviations to use to define the decision boundary are present in the literature (e.g., compare 12, 29, 37).

### 5.1 Carlos Dopico *et al., 2021*

We used the dataset available from Castro Dopico et al. (7), with the group variable representing the batch divisions. This dataset comprises the log-transformed normalised OD for IgG responses against stabilized trimers of the SARS-CoV-2 spike glycoprotein (SPIKE) and the smaller receptor-binding domain (RBD) in 2,100 sera samples from blood donors, 2,000 samples from pregnant volunteers, 595 historical negative controls, repeatedly sampled, and 149 PCR-positive patients (positive controls from 8). The data were generated across seven batches, with the positive controls contained in two of these. This, combined with our expectation that seropositivity should increase with time as more of the population were exposed to SARS-CoV-2, suggests that the batch and seropositivity frequency are dependent. Based on our simulation study, we would expect that a pre-processing batch normalisation would therefore produce misleading results.

We ran five chains of the MVT mixture model for 50,000 iterations for each of nine combinations of different choices for the hyperparameters of the batch effects in the model (choices in table 1, distributions in figure 4 A and B). The first 20,000 samples were removed as burn-in, and we thinned to every 100th sample to reduce auto-correlation.

**Table 1:** Hyperparameter combinations used in analysing the data from Castro Dopico et al. (7). The prior expected value of the batch scaling effect is the same for all choices of *α* and *β*. The choice of λ represents the scale we *a priori* expect for the batch shift effect.

|  | Value |  |  |  |  |  |  |  |  |
| --- | --- | --- | --- | --- | --- | --- | --- | --- | --- |
| $\alpha$ | 1 | 5 | 10 | 1 | 5 | 10 | 1 | 5 | 10 |
| $\beta$ | 3 | 11 | 21 | 3 | 11 | 21 | 3 | 11 | 21 |
| $\lambda$ | 0.01 | 0.01 | 0.01 | 0.10 | 0.10 | 0.10 | 1.00 | 1.00 | 1.00 |

**Figure 4:**
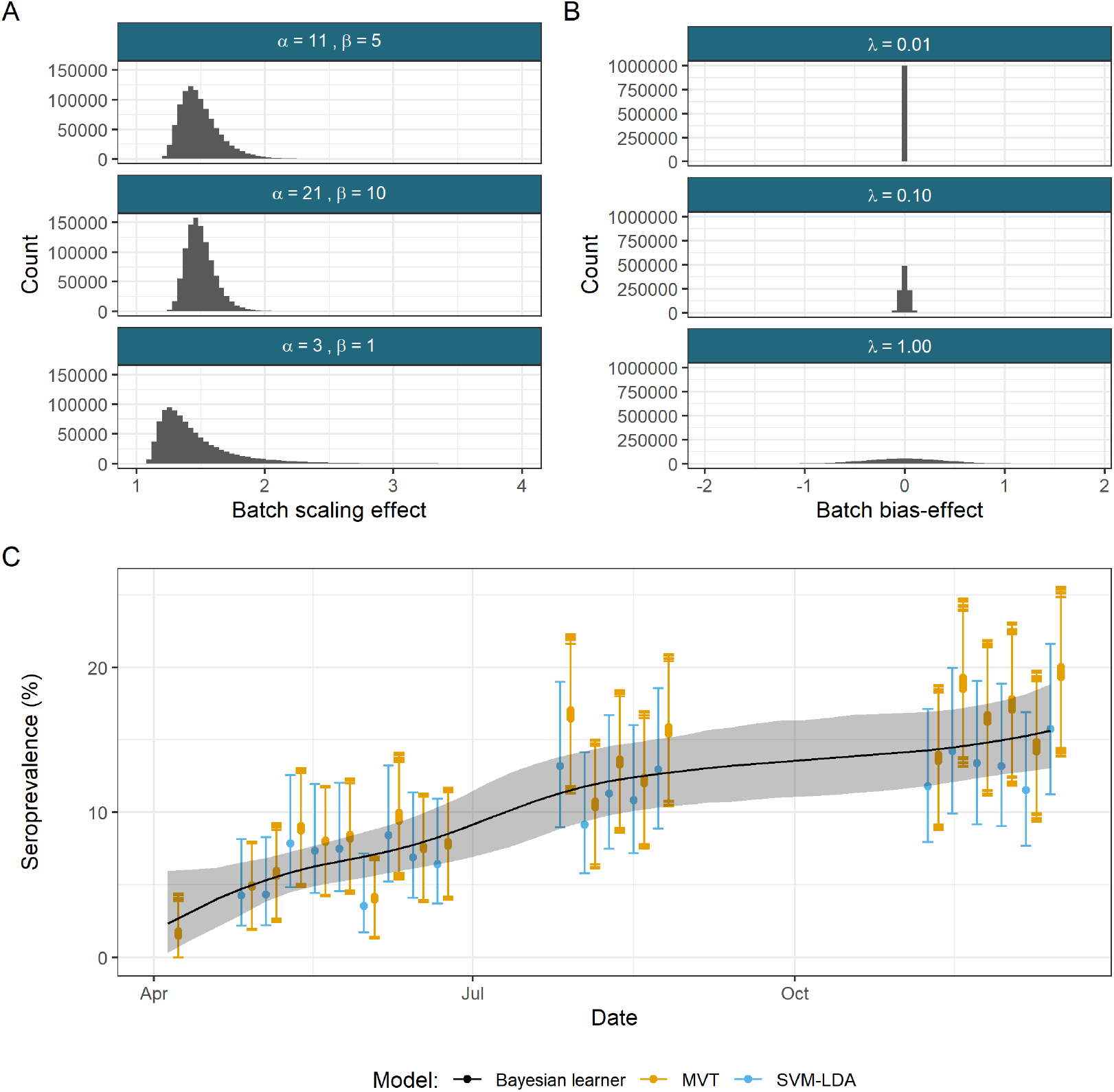
Effect of hyperparameter choices on seroprevalence estimates. One million draws from the prior distributions for the different hyperparameter choices for A) the batch scaling effect and B) the batch shift effect. In A) draws exceeding a value of 4 are hidden. This means that approximately 0.5% of the draws from the prior distribution with a shape of 3 and a scale of 1 are not shown. C) A comparison of the estimated seroprevlaence with population 95% confidence intervals for the MVT mixture model with nine different choices of hyperparameters for the batch-effect prior distributions and the estimates from Castro Dopico et al. (7) for the SVM-LDA ensemble model and the Bayesian learner from Christian and Murrell (9). The Bayesian learner is designed to estimate seroprevalence during an epidemic and provides a smooth, nondecreasing estimate across time. Its assumptions ensure a more consistent increase across time, whereas the SVM-LDA and MVT mixture models are not incorporating any explicit temporal information. The estimates from the mixture model have been moved 3 days to the right on the x-axis to reduce overlap.

We chose a representative chain for each hyperparameter combination to estimate the seroprevalence for each week of the year 2020 for which samples are available and compared these to the estimates from Castro Dopico et al. (7) (figure 4 C). Our point estimate was the mean posterior probability of allocation for the non-control data. This was highly consistent across hyperparameter choices and was contained within the confidence interval of the estimate provided by Castro Dopico et al. (7). However, our seroprevalence point estimates, particularly in later dates, were higher than the those from Castro Dopico et al. (7). Table 1 of the Supplementary material shows the point estimate from the ML methods used in the Simulation study, our MVT mixture model and that from the original paper. This shows that while our method provides higher point estimates than those from Castro Dopico et al. (7), the other ML methods (barring the SVM) provide estimates much closer to or even exceeding that from the MVT.

The seroprevalence estimates and their credible intervals were almost identical across hyperparameter choices, suggesting that the classification results are robust to different choices for these hyperparameters. We took a single chain with hyperparameter choice *α* = 5, *β* = 11 and λ = 0.1 as a representative example. This value of λ represents our expectation that *m_b_* should be approximately an order of magnitude smaller than *μ_k_*. We used this to infer a point classification and a batch-corrected dataset (figure 5). Note that the data were on a similar scale to the observed data (figure 5), the lack of identifiablity for parameters in the likelihood function did not emerge as a problem here. The batch-corrected dataset was better visually separated into seronegative and seropositive classes than the observed data due to our batch-correction.

**Figure 5:**
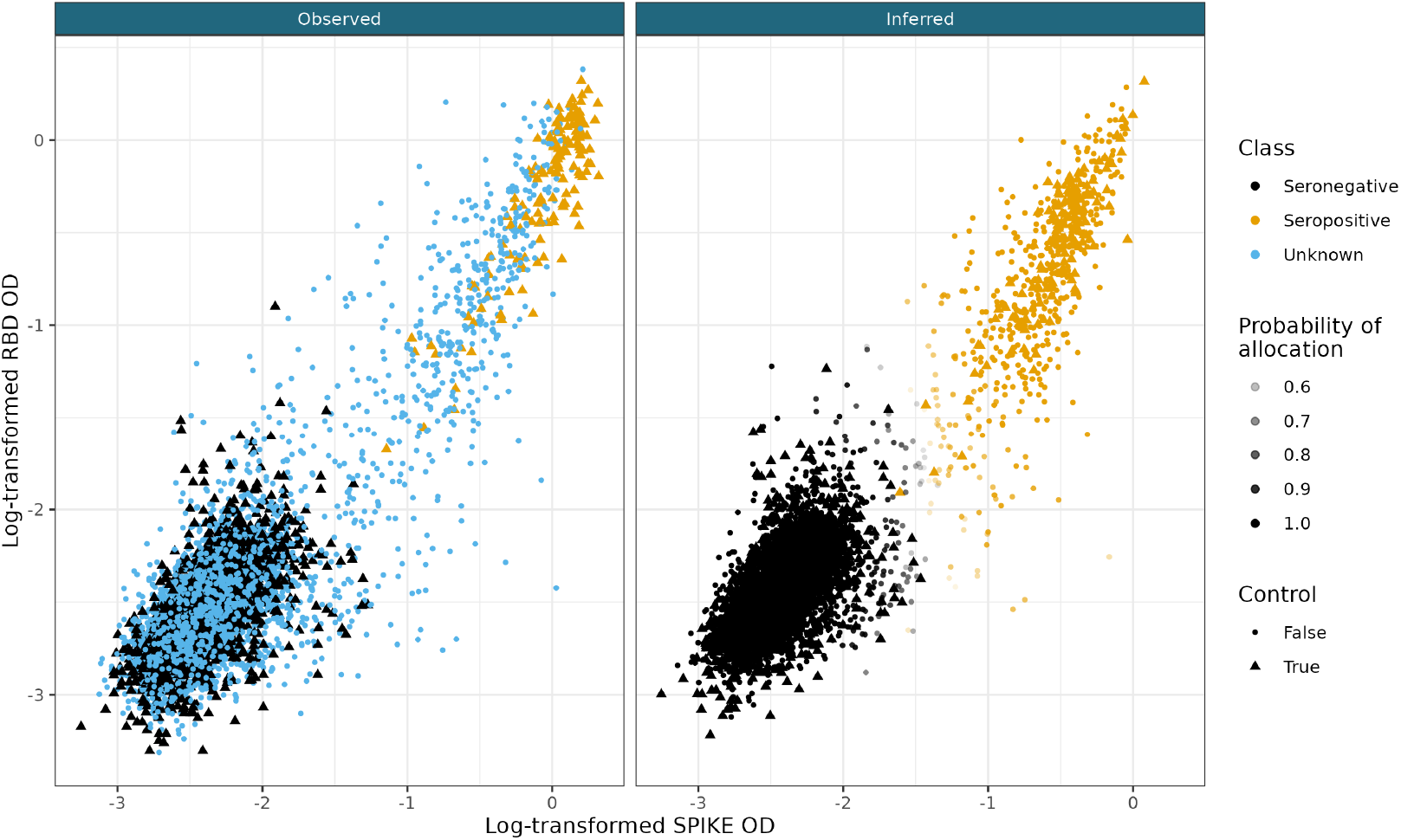
In the first facet, the observed data from Castro Dopico et al. (7) and on the right, the point estimate of the inferred, batch-corrected dataset from the MVT mixture model with *a* = 11, *β* = 5, λ = 0.1. Points on both plots are coloured by the class. In the observed dataset non-control points are labelled “Unknown” and in the batch-corrected dataset these points are labelled with their inferred class and have their opacity controlled by the inferred allocation probability.

To confirm the batch-correction was working as intended we considered repeated control samples from a particular patient, “Patient 4”, and the negative controls in batches with a high proportion of negative controls (> 100). The Patient 4 samples were all collected at the same time and were included in several plates as a positive control but discarded before our analysis because it was chosen for extremely high antibody levels and so is unrepresentative, even for the seropositive class. We hypothesised that appropriate batch-correction should bring the different measurements of this sample closer together, which is indeed what we observed after applying the correction learnt from the samples excluding Patient 4 (figure 6 A). Before correction, the batches had no overlap; there was a distance of 0.197 between the batch means. After correction the two batches overlapped with a distance of 0.040 between the means as the points moved closer together and towards the class mean (figure 6 A would correspond to the upper right hand of figure 5). For the set of negative controls we consider, we expected that these batches should have overlap more after correction as their relationship between batch and class should be similar. This is what we observed (figure 6 B) and the variance explained in the SPIKE and RBD optical density by the batch variable for these samples is significantly lower after the batch-correction. For SPIKE OD, 4.2% of the variation is explained by batch of origin in the observed data, 0.5% is explained in the batch-corrected data. For the RBD optical density, the respective figures are 4.3% and 1.4%.

**Figure 6:**
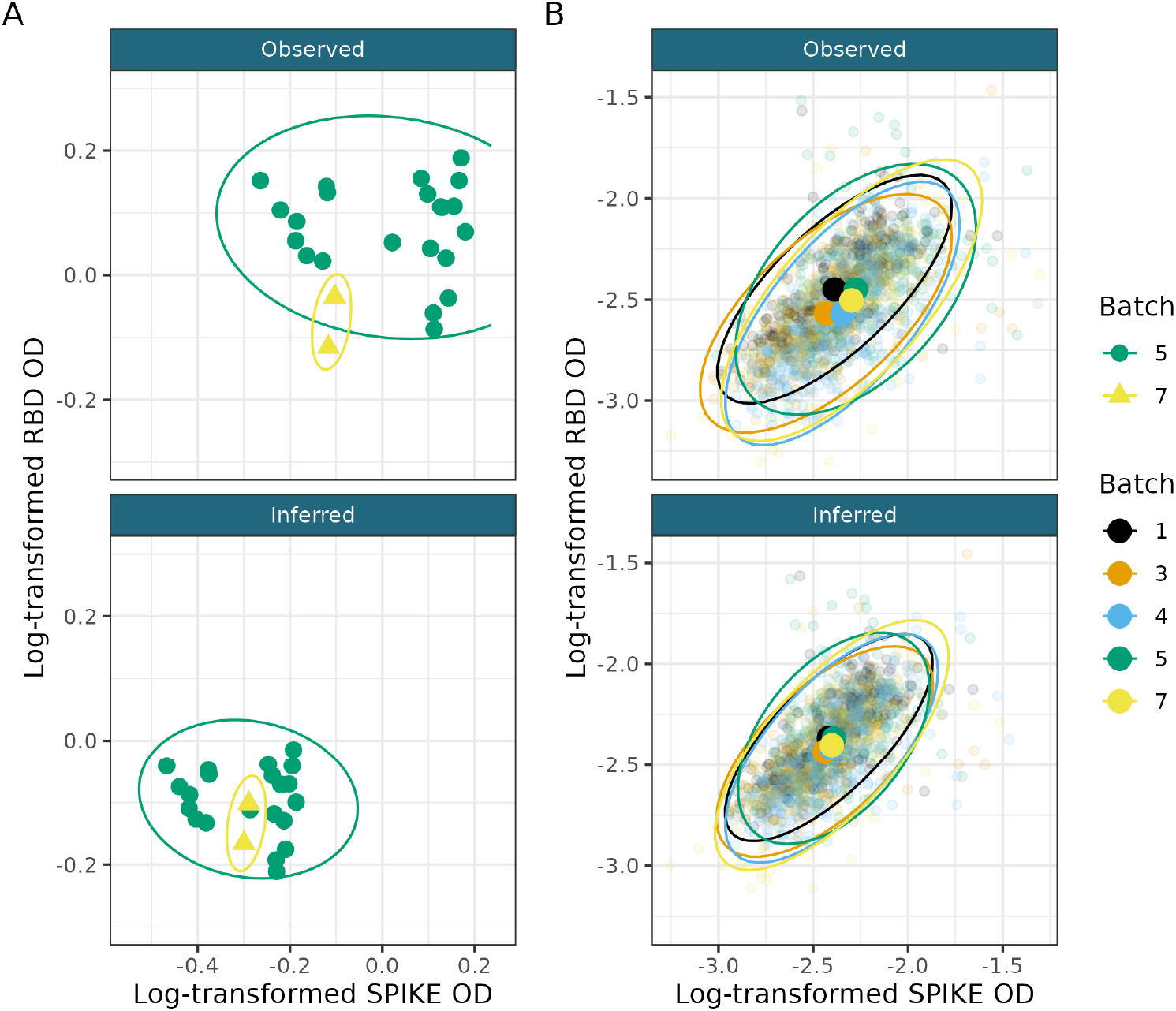
A) The samples from Patient 4 as observed and after batch correction. B) The mean and covariance for the batches containing a large number (> 100) seronegative controls as observed and after batch-correction. Note that the means overlap significantly more and relationship between SPIKE and RBD is more uniform across batches after the correciton.

### 5.2 Pseudo-ELISA data

We wished to investigate the possibility that other known positive samples could be more extreme than the non-hospitalised donors. To examine this, we generated datasets from the model fitted in section 5.1. This also tests if the model has learnt representative parameters for the dataset, as our generated data should be very similar to the original data. We used the MCMC sample mean for each parameter except the class weights. For the class weights we used the inferred proportion of each class in each batch to preserve the problem of the imbalance of classes across batches. In the original data, the positive controls were more extreme members of the positive class, having sufficiently severe symptoms to have undergone PCR testing when such resources were severely constrained early in the pandemic. To reflect this in our data generation procedure, we increased the probability that samples with observed positive labels (i.e., the positive controls) are from the tail of the distribution of the seropositive measurements which is furthest from the seronegative class, whereas the negative controls are sampled uniformly from the seronegative population. An example dataset is shown in figure 7 C, note how closely it resembles the true ELISA data in figure 5, suggesting that the model has learnt accurate values. See section 8 of the supplementary material for a deeper explanation of the generation process.

**Figure 7:**
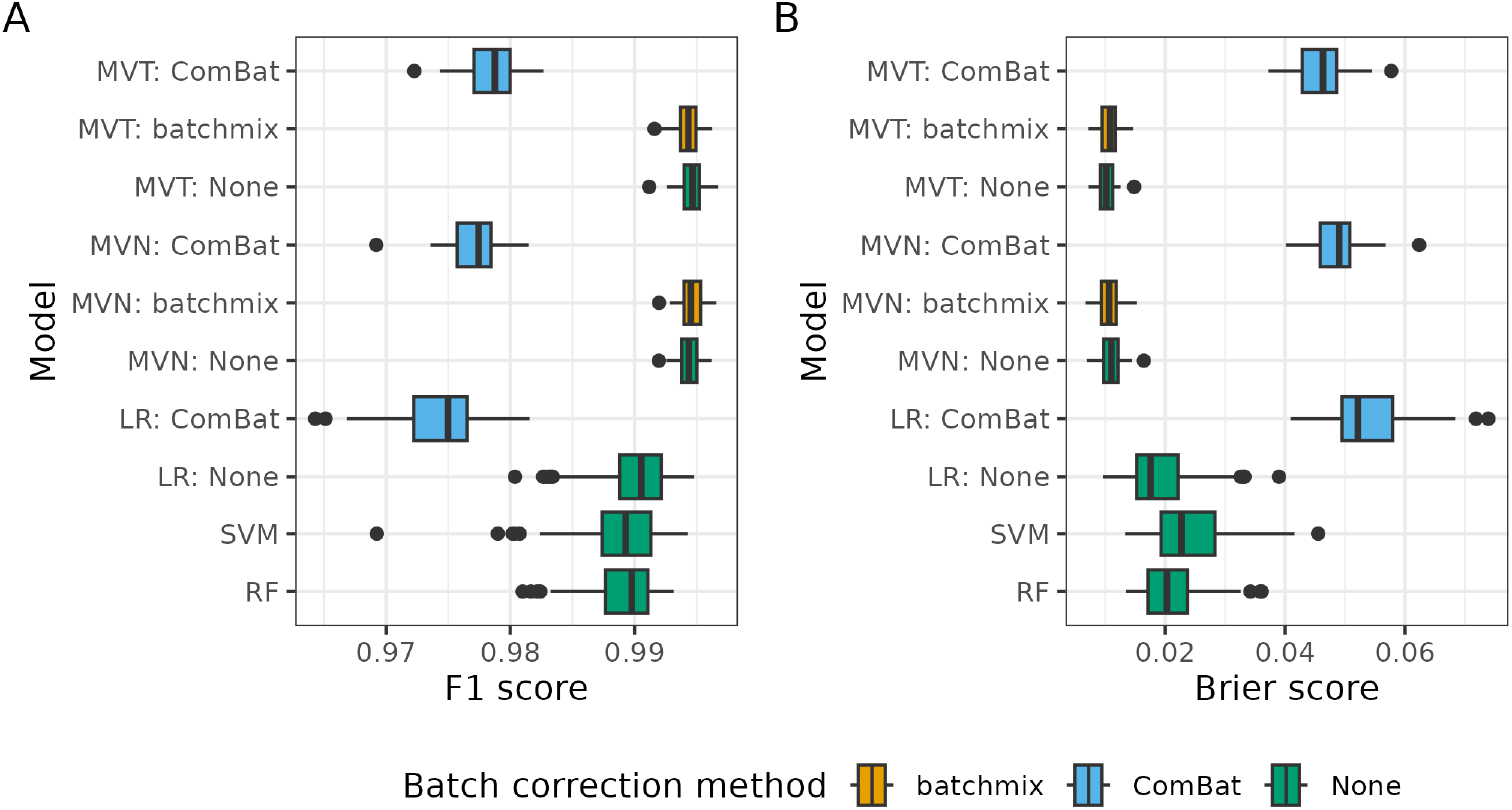
Performance of different methods across 100 datasets simulated from the converged MVT model for the ELISA data from Castro Dopico et al. under A) the F1 score and B) the Brier score.

We performed a similar analysis to our original simulation study on these datasets, comparing our models to a range of off-the-shelf machine learning methods. Across all of the simulations, we found that our mixture models outperformed other methods under both the F1 score and the squared distance (figures 7 A, 7 B). The semi-supervised Bayesian mixture model also performed well here; this is due to the large proportion of negative samples in a single batch and the large imbalance in class sizes (approximately 19 to 1).

### 5.3 Dingens et al., 2020

As a final real data example, we analysed the ELISA data collected by Dingens et al. (12). This consisted of 1,891 measurements of antibodies to the SARS-CoV-2 RBD protein. 1,783 of these were from residual serum from Seattle Children’s Hospital, with 52 pre-2020 samples used as negative controls and 52 samples from individuals with RT-PCR-confirmed infections as positive controls (figure 8 A). These data are different to the data from Castro Dopico et al. (7) in several ways. There is only a single antigen, there is a smaller ratio of controls to non-controls, particularly for the seronegative samples, and the controls do not appear to be representative of either class. The mean log OD of the negative controls is −1.91, whilst the dataset mean is −2.28 without controls. We analyse the log-transform of the OD using our MVT model for the same variety range of hyperparameter choices as in table 1. An example of a batch-corrected dataset is shown in figure 8 B. We show the comparison of the inferred seroprevalence in each batch for an example chain of each of these models as well as that estimated by Dingens et al. (12) (figure 8 C). The 9 different hyperparameter choices have almost identical seroprevalence estimates and are estimating higher levels of seroprevalence than the estimate provided by Dingens et al. (12).

**Figure 8:**
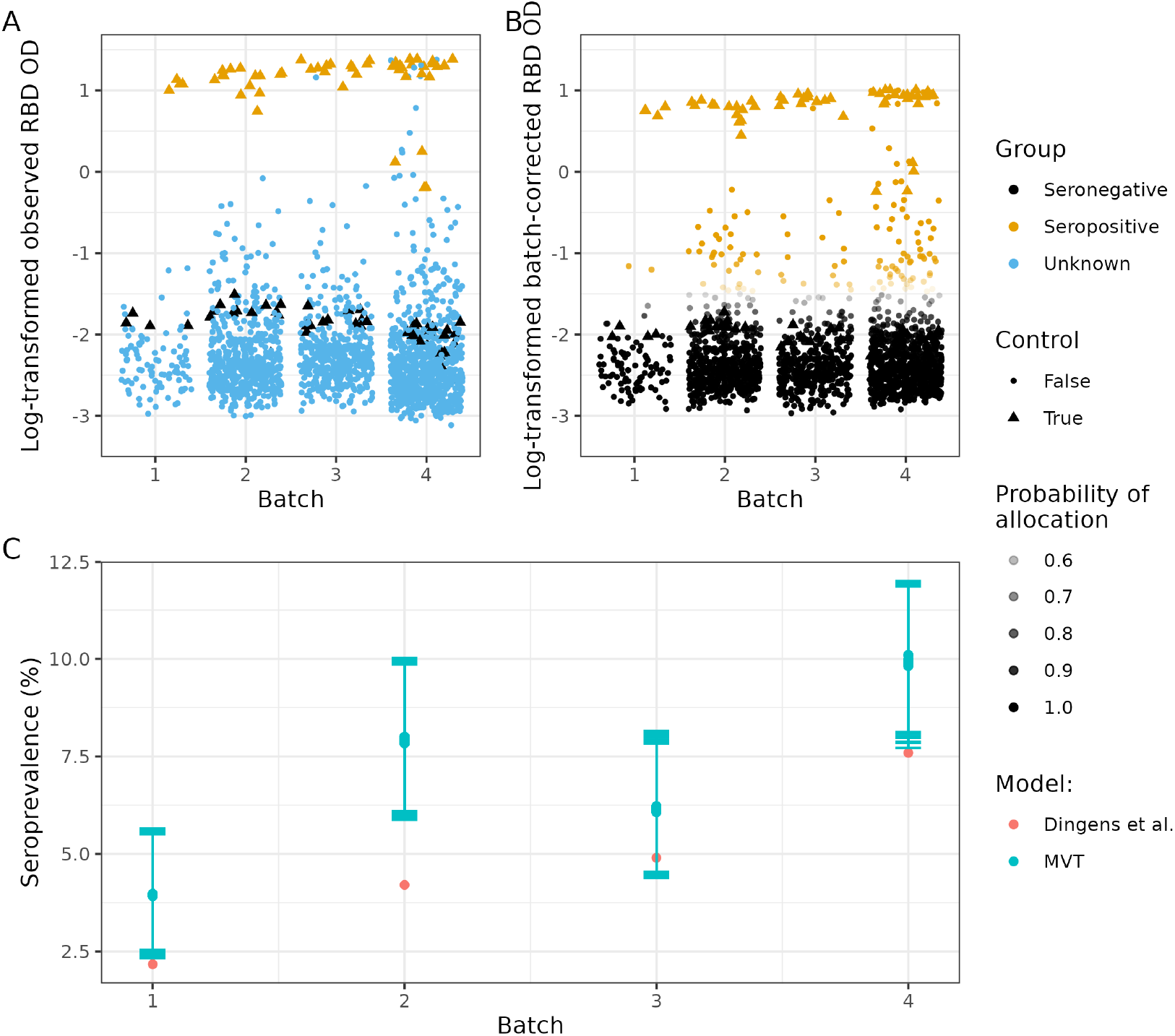
A) The observed data from Dingens et al. (12) and B) the point estimate of the batch-corrected dataset from the MVT mixture model with *α* = 11, *β* = 5, λ = 0.1. Points on both plots are coloured by the class. In the observed dataset non-control points are labelled “Unknown” and in the batch-corrected dataset these points are labelled with their inferred class. C) A comparison of the seroprevalence estimate from the MVT mixture model with nine different choices of batch-effect hyperparameters and that from Dingens et al. (12). The error bars indicate the 95% credible interval for the seroprevalence estimates of the MVT mixture model in each batch; this is not available for the estimate from Dingens et al. (12).

## 6 Discussion

The results of our simulation study show that our mixture model consistently matches or outperforms several alternatives when applied to data with batch effects, across a range of data generating models. We believe that the simulation study and the gene expression analysis shows our method is a good model choice in a low-dimensional setting. If batch and class are independent, it performs almost identically to ComBat, but it is much stronger when this dependency exists. On the other side of the spectrum, using the standalone mixture model with no batch-correction is also a viable option, but when batch effects are large (as in the Varying batch effects scenario), this is significantly worse than including a batch correction. As the relationship between batch and class and the magnitude of the batch effects is rarely known, we believe our model offers a better guarantee of meaningful inference. In the more specific scenario where data were generated from a converged chain that had been applied to the ELISA data from Castro Dopico et al. (7), we obtained the same findings, with our model again performing better than the off-the-shelf machine learning methods and ComBat over-correcting for batch due to the imbalance of classes. We also see from our simulation study that we should use the MVT density over the MVN density, as the MVT can approximate the MVN quite well by learning a large degree of freedom, but also has additional flexibility as shown by the Multivariate t generated simulation scenario where the MVN mixture model behaved very inconsistently. The only cost of the MVT mixture model is the approximate 50% increase in runtime, but as our implementation is quite fast we believe that this is not a significant detractor. Based on these results we recommend the use of our MVT mixture model when the analyst suspects the classes in the data may be non-Gaussian.

In terms of estimating seroprevalence, our mixture model performed very well in our simulation study. Using the results shown in figure 1 B, we can try to gauge how well our method is performing in the ELISA data. We would argue that the most pertinent scenarios are the MVT generated (the ELISA data are non-Gaussian), the Varying batch effects and the Varying class representation scenarios. Our method estimates seroprevalence close to the truth, or slightly smaller, in these simulations. Based on this, we suspect that the high estimates of seroprevalence provided by our model (relative to those from the original papers) in the ELISA analyses are plausible.

In the Swedish dataset, we are reassured that the batch-correction is reasonable by our analysis of the patient 4 samples - these samples were used across several batches as positive controls; after applying the correction learnt on the dataset excluding these extreme samples they are no longer separable by batch and have moved towards the class mean. The data generated from our converged model also appears very similar to the observed data, suggesting that the model assumptions are reasonable, and that meaningful estimates of the parameters were obtained.

In the analysis using the data from Dingens et al. (12), the unrepresentative negative controls presented a problem. We believe that the preceding analyses show the potential advantages of our model over existing methods, but this dataset is a good example to show that our method is not a panacea that may overcome all problems - it remains vital to have useful and relevant data in order to perform meaningful inference (13). Any analysis that uses training data that appear to be drawn from a different population than the test data is unlikely to produce meaningful results. Furthermore, the data are not well-described by a pair of MVT distributions (even allowing for our additional flexibility with the batch parameters). This combination of model misspecification and misleading training data makes us skeptical of the inferred parameters.

We note, however, that in the simulation of pseudo-ELISA data, our method still performed strongly despite the positive controls not being representative of the general seropositive sample. In this case our model was correctly specified (the data are generated from a MVT mixture model). In general, we suspect that our method is useful if either the assumption that the labelled data represent their class well or that the model density choice is correct are slightly relaxed, but if both do not hold or if either is profoundly wrong then the model will perform poorly.

Since only the combined class and batch parameters, *μ_k_* + *m_b_* and Σ_*k*_ ⊕ *S_b_*, are identifiable, one might expect this to present challenges when fitting our model. However, Redner (35) showed that the maximum-likelihood estimator is consistent when the distributions are not identifiable and the nonuniqueness is caused by the particular parameterization. Note that even if the individual batch and class parameters never stabilise (note that their combinations are identifiable and should converge), running multiple chains helps to avoid this potential pitfall as one could use the trace plots for the complete likelihood to assess if the chains have reached a common mode in the likelihood surface even if the individual batch and class parameters did not converge. This is standard practice when using stochastic methods, so this aspect of the model should not introduce additional work to the recommended Bayesian workflow (15). Furthermore, from the similarity of the inferred parameters across multiple chains in the Base case simulation (figure 2), we have empirical evidence that this behaviour is not common. We also saw that the seroprevalence estimates and their credible intervals across different hyperparameter choices in the ELISA analyses were well-behaved and, as a result, so was the inferred allocation. This similarity across hyperparameter choice suggests that choosing between specific values is not too important, but we suspect that, if the sample size is smaller, having λ close to one could exacerbate the identifiability problem for the batch shift effect and the class mean. Therefore, we suggest setting λ ≤ 0.1 to encourage these parameters to converge in the small sample setting (although note that their sum, *μ_k_* + *m_b_*, should converge regardless).

We have developed a Bayesian method to predict class membership and perform batch-correction simultaneously, developing on the pre-processing, univariate method of Johnson et al. (19). Our method is intended for low-dimensional data, but the main limitation for higher dimensional data is computational (inverting the covariance matrix becomes very costly) rather than theoretical. Our model is not strictly limited to the semi-supervised setting either; it could be used for unsupervised learning. In this case we expect that the model will rely much more heavily on the distributional assumptions. Our work could be extended to include alternative densities, such as the skew multivariate t We could extend the model to include batch-specific class weights, such as we used to generate the data in our Varying class representation simulation scenario, or a deeper hierarchy for the batch parameters, such as nested batches (e.g., this could represent scenarios where multiple plates are run at each of multiple time points or locations).

## Acknowledgements

We would like to thank Dingens et al. (12), specifically Janet A. Englund and Jesse D. Bloom, for being willing to openly share their data and batch information.

## Authors’ contributions

SC, PK and CW all contributed to model design. SC implemented the model in C++ and built the R package with PK and CW contributing to debugging strategies. SC designed the simulation study and the pseudo-ELISA data. SC, PK and CW all contributed to the design of the Metropolis algorithm used to implement the model and the choice of proposal densities. PK, CW and SC all contributed to analysis and the interpretation of results. KN wrangled and cleaned the gene expression data. XD and GK generated the first ELISA dataset which CW cleaned. All authors read and approved the manuscript.

## Supplementary material

### Abstract

Description of the model, our choice of priors, and the sampling algorithm. Example of likelihood trace plots for model convergence. Description of how the simulated data is generated for both the main simulation study and the pseudo-ELISA simulation.

### 1 Model

Our data *X* = (*X*_1_, …, *X_N_*) is generated across *B* batches where the origin batch of each point is known and represented by the vector *b* = [*b*_1_, …, *b_N_*]^⊤^. We are interested in classifying *X* into *K* disjoint classes. We model *X* using a *K* component mixture model:

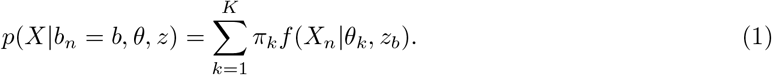

Here *f* (·) is the density function, π = [π_1_, …, π_*k*_]^⊤^ are the component or class weights, *θ* = (*θ*_1_, …, *θ_k_*) are the parameters describing the classes and *z* = (*z*_1_, …, *z_B_*) are the parameters associated with the batches. We introduce an allocation variable, *c* = [*c*_1_, …, *c_N_*]^⊤^, to represent the class membership and assume that each class is represented by a single component of the mixture. Conditioning on *c*, our model is then

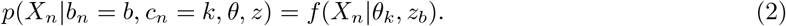

In our motivating example, *c* contains some observed values (alternatively, *c* contains missing values), this enables supervised or semi-supervised methods to infer the missing values. We introduce a binary vector, *ϕ* = [*ϕ*_1_, …, *ϕ_N_*]^⊤^, indicating if the label of the *n^th^* individual is observed or not. If we separate our dataset into subsets of labelled and unlabelled data

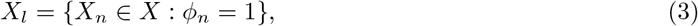

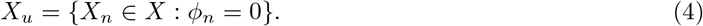

and use *X_l_* to train some classifier which predicts the labels of *X_u_* (as we do with the off-the-shelf ML models in our simulation study), our method would be a supervised classifier. However, the Bayesian framework enables us to integrate these steps in a semi-supervised model, using the inferred allocations to include *X_u_* in modelling the class and batch parameters.

#### 1.1 Multivariate Normal

Let *f* be the density function for the multivariate normal distribution, parametrised by a mean vector *μ* and a covariance matrix Σ.

We assume

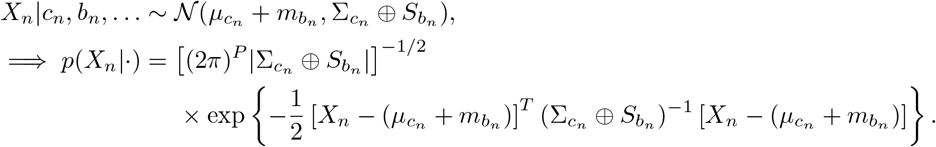

We also assume that the batch effects have no correlation across dimensions. We restrict the covariance matrix, *S_b_*, to being diagonal and assume independence between the entries of *m_b_*.

Our hierarchical model is

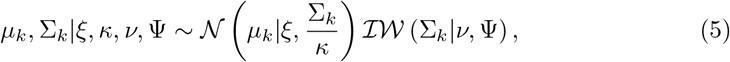

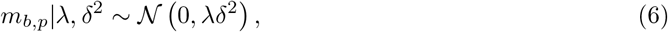

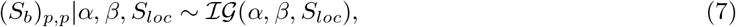

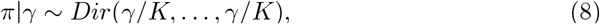

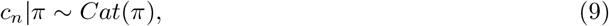

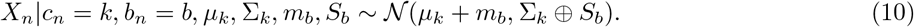

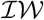 denotes the inverse-Wishart distribution, 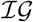 denotes the inverse-Gamma distribution with a shape *α*, rate *β* and location *S_loc_*. 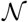 is the Gaussian distribution, *Dir* is the Dirichlet distribution and *Cat* is the categorical distribution. As we assume independence of batch effects across dimensions, we model each entry of the *b^th^* batch mean vector, *m_b,p_*, and the *b^th^* batch covariance matrix, (*S_b_*)_*p,p*_, using one dimensional distributions.

**Figure 1:**
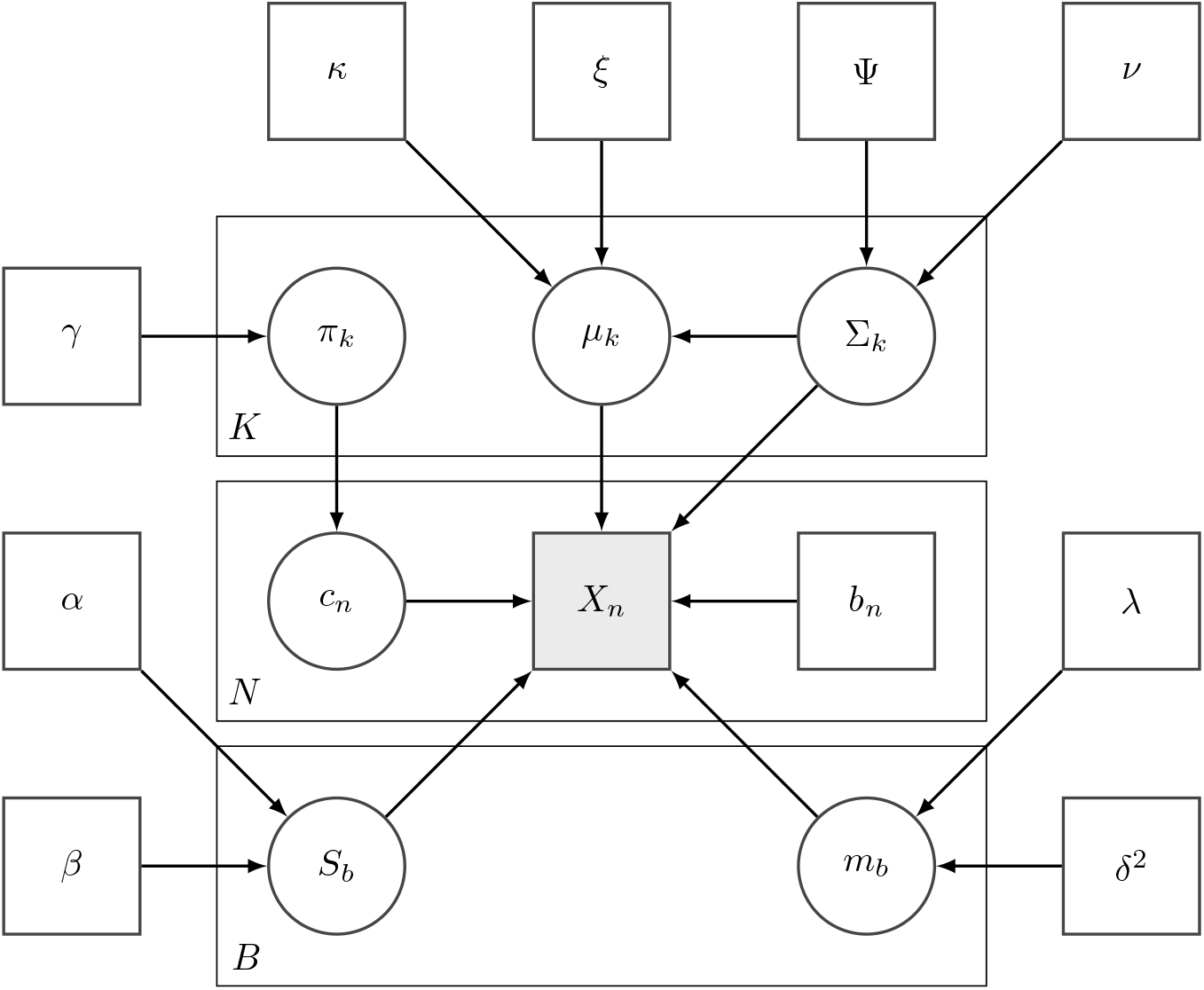
Directed acyclic graph for mixture of multivariate normal distributions with random effects. Squares indicate observed or known quantities. Note that a subset of *c* is observed in our application.

The total joint probability is

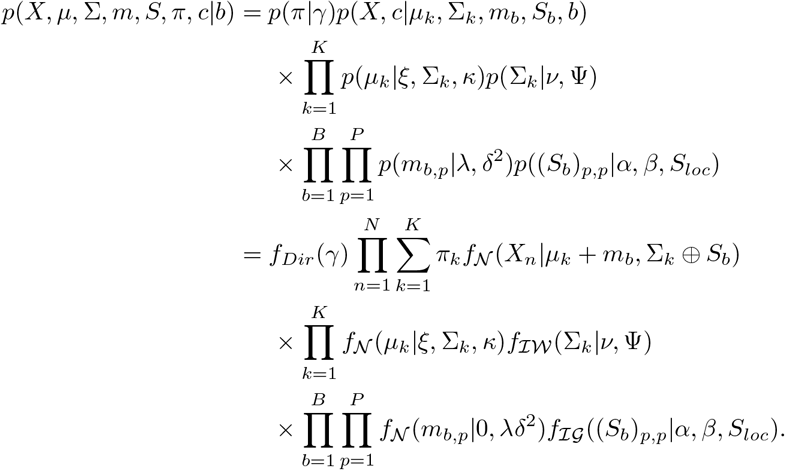

#### 1.2 Multivariate t

If we let *f* be the density function for the multivariate *t* (**MVT**) distribution, parametrised by a mean vector *μ*, a covariance matrix Σ and degrees of freedom, *η*, then the model remains as described in section 1.1 and equations 5, except the model likelihood changes and we introduce a prior distribution over *η*:

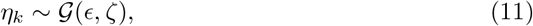

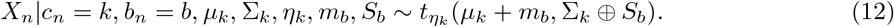

here 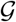 denotes the Gamma distribution parametrised by a shape and rate.

The total joint probability for the mixture of MVT distributions is

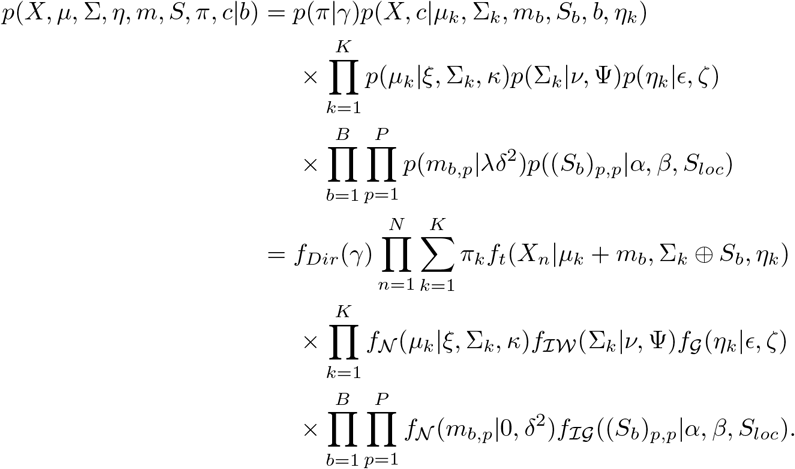

#### 1.3 Parameter interpretation

Note that the “batch” parameters should not be inferred as direct estimates of the effect the batches have on the true measures. As we are essentially performing a classification on the inferred batch-free dataset,

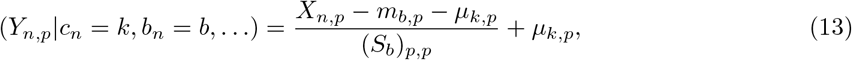

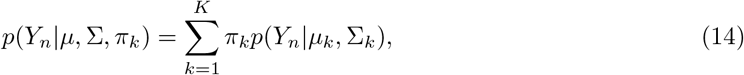

and the likelihood parameters of *μ_k_* + *m_b_* and Σ_*k*_ ⊕ *S_b_* are not constrained in the likelihood, we recommend that users focus on the relative change in the measurements for batches, the inferred dataset and the inferred classification rather than the direct meaning of individual parameters.

### 2 Empirical Bayes

We use the suggestions of Fraley and Raftery (2007) for our choices of prior hyperparameters on the class parameters.

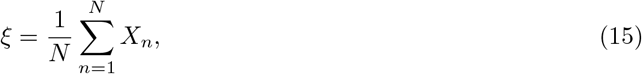

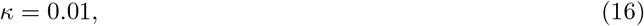

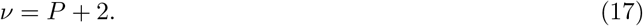

The choice of *ξ* is self-explanatory. *κ* can be viewed as the number of observations contributing to the prior. Fraley and Raftery (2007) choose a value based on experiments to acquire a BIC curve that is a smooth extension of the counterpart without a prior. The marginal prior distribution of *μ_k_* is a Student’s *t* distirbution centred at *ξ* with *ν* – *P* +1 degrees of freedom. *ν* is the smallest integer value for the degrees of freedom that gives a finite variance.

We set Ψ as a diagonal matrix. Let

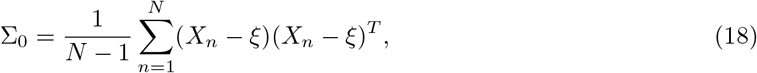

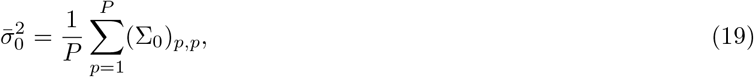

then

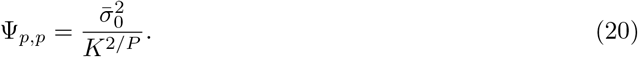

The logic is that the mixture components are expected, *a priori*, to each fill a common fraction of the total volume of space the data occupies.

For the concentration on the class weights, we use a flat prior with *γ* =1. In our motivating exmple of ELISA data, we cannot use more information (such as the ratio of class members in the known data), as the negative controls are historical samples the number of which is chosen before the experiment and is not related to the expected seroprevalence in the dataset.

For the degrees of freedom for the MVT, *η_k_*, we use an uniformative prior that offers a range of plausible values, *ϵ* = 2.0, *ζ* = 0.1 (Juárez and Steel, 2010).

### 3 Sampling algorithm

We use a *Metropolis-within-Gibbs* algorithm to sample our parameters. All parameters where the form of their posterior distribution is known are sampled via Gibbs sampling (Geman and Geman, 1984), the remaining parameters are sampled in a Metropolis-Hastings step (Metropolis et al., 1953; Hastings, 1970).

#### Algorithm 1: *sampler*(*X*, *I*, *c*_0_, *fixed, b, K*)

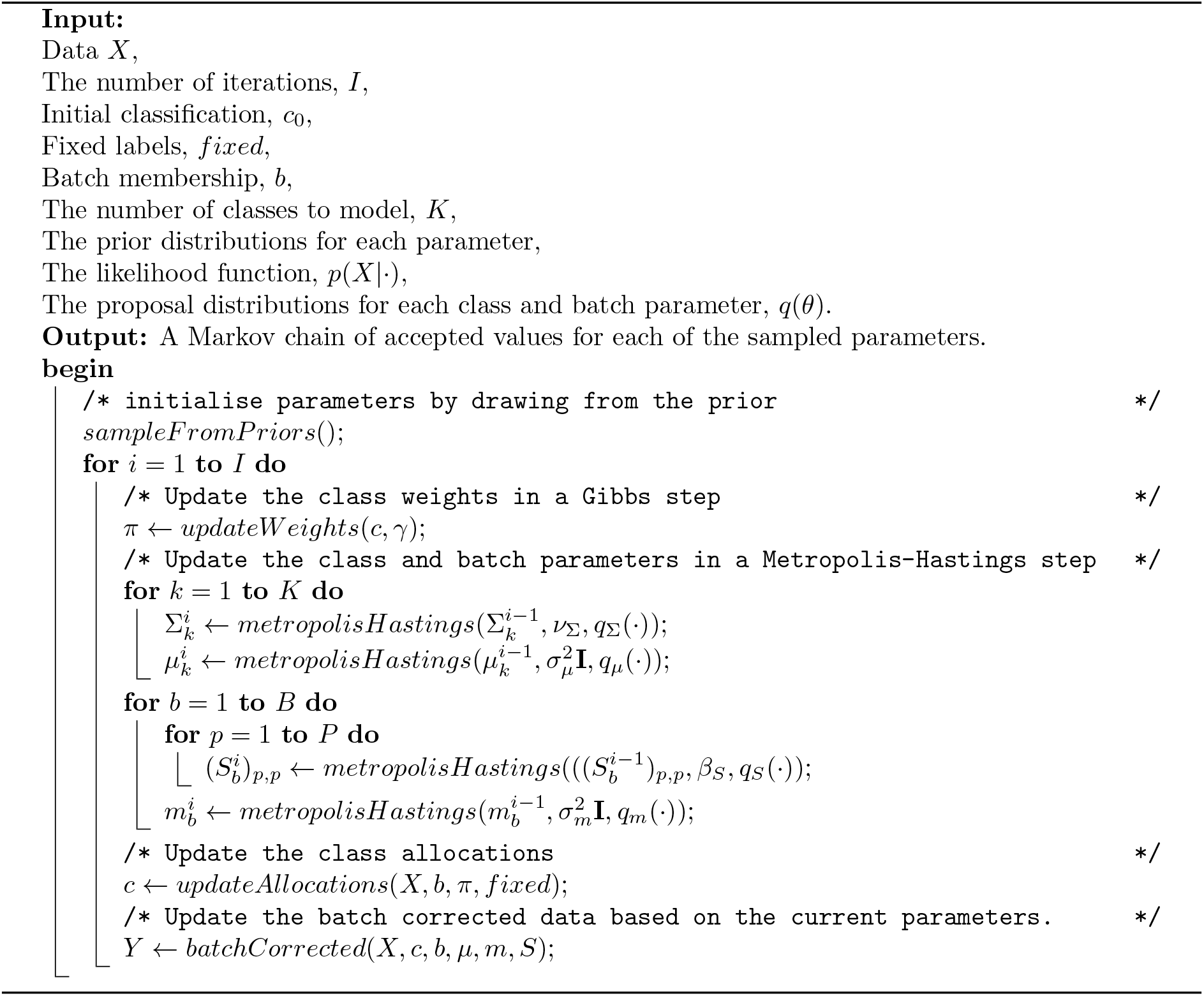

#### 3.1 Proposal distributions

For our batch and class parameters, we choose proposal densities that have an expectation of the current value and have the correct support. The class and batch means have a support (∞, ∞); this allows use of a Gaussian proposal distribution with a mean of the current value.

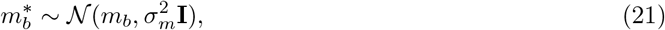

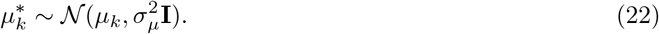

##### Algorithm 2: sampleFromPriors()

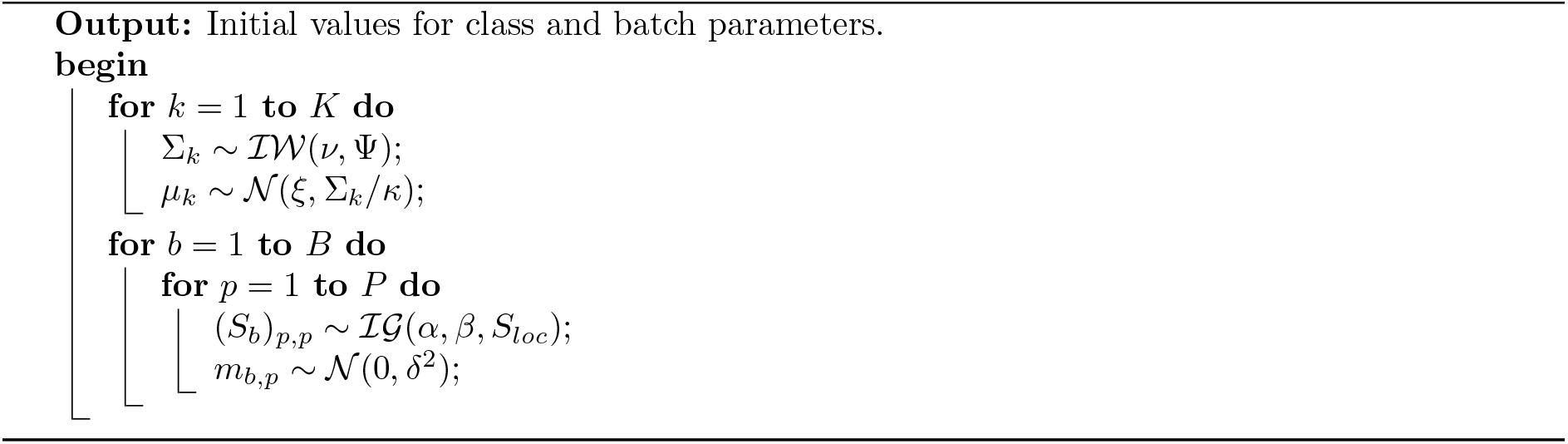

##### Algorithm 3: updateAllocation(X, b, π, fixed)

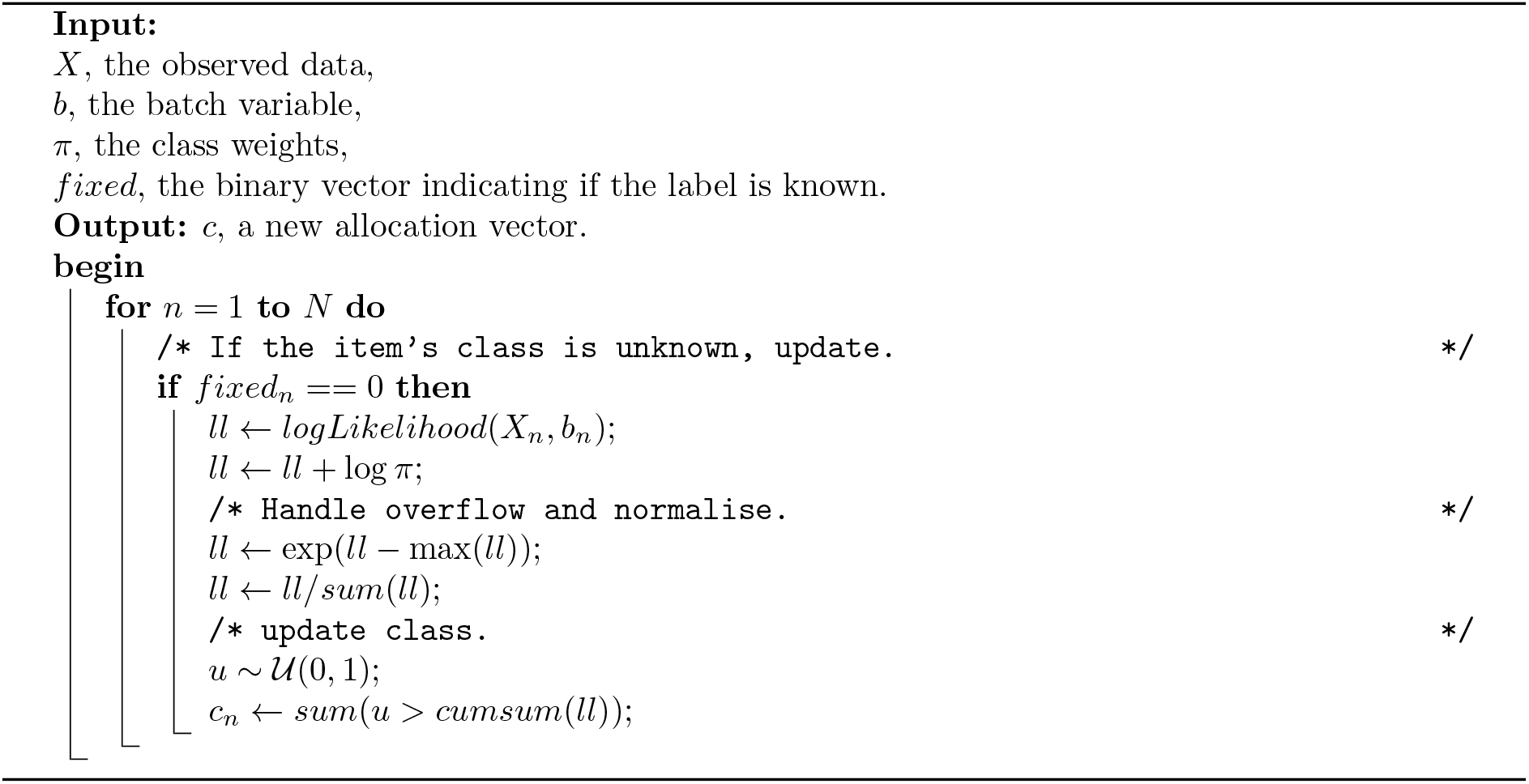

##### Algorithm 4: updateWeights(c, γ)

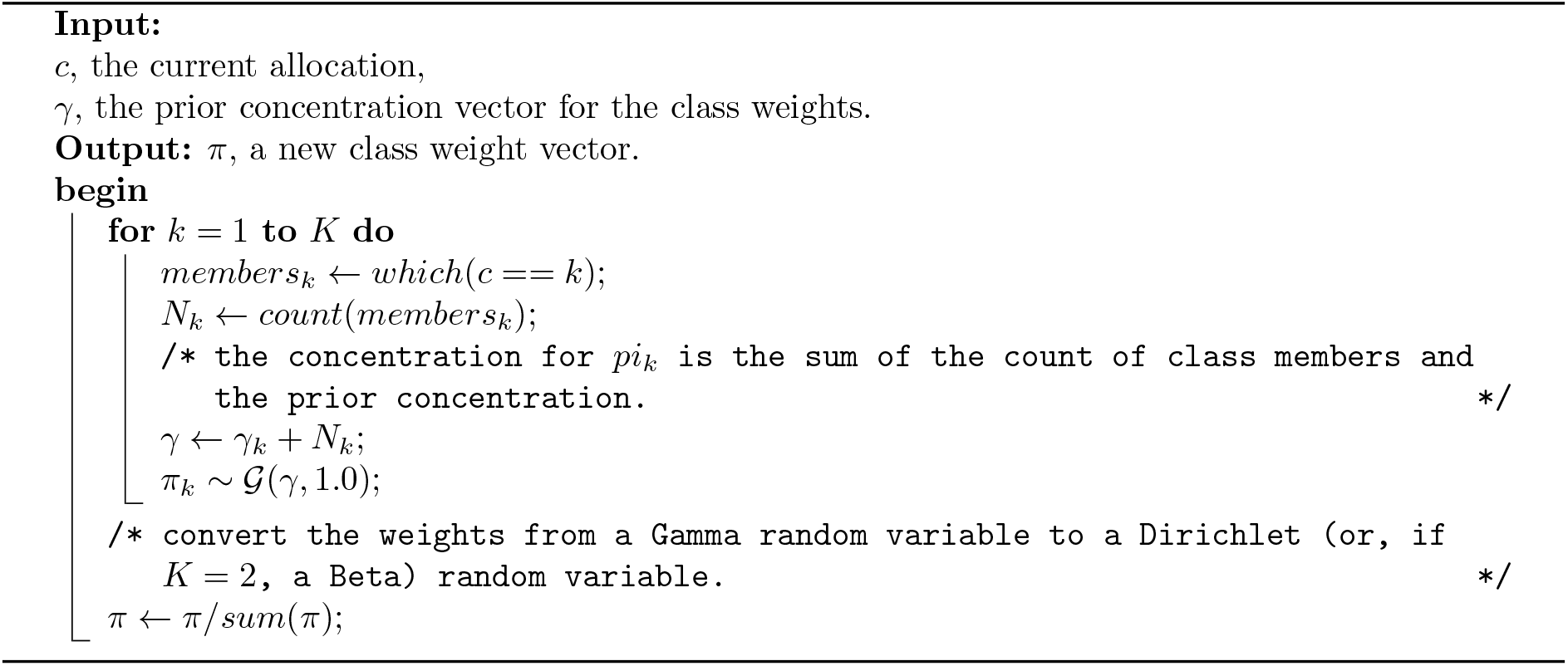

##### Algorithm 5: batchCorrected(X, c, b, μ, m, S)

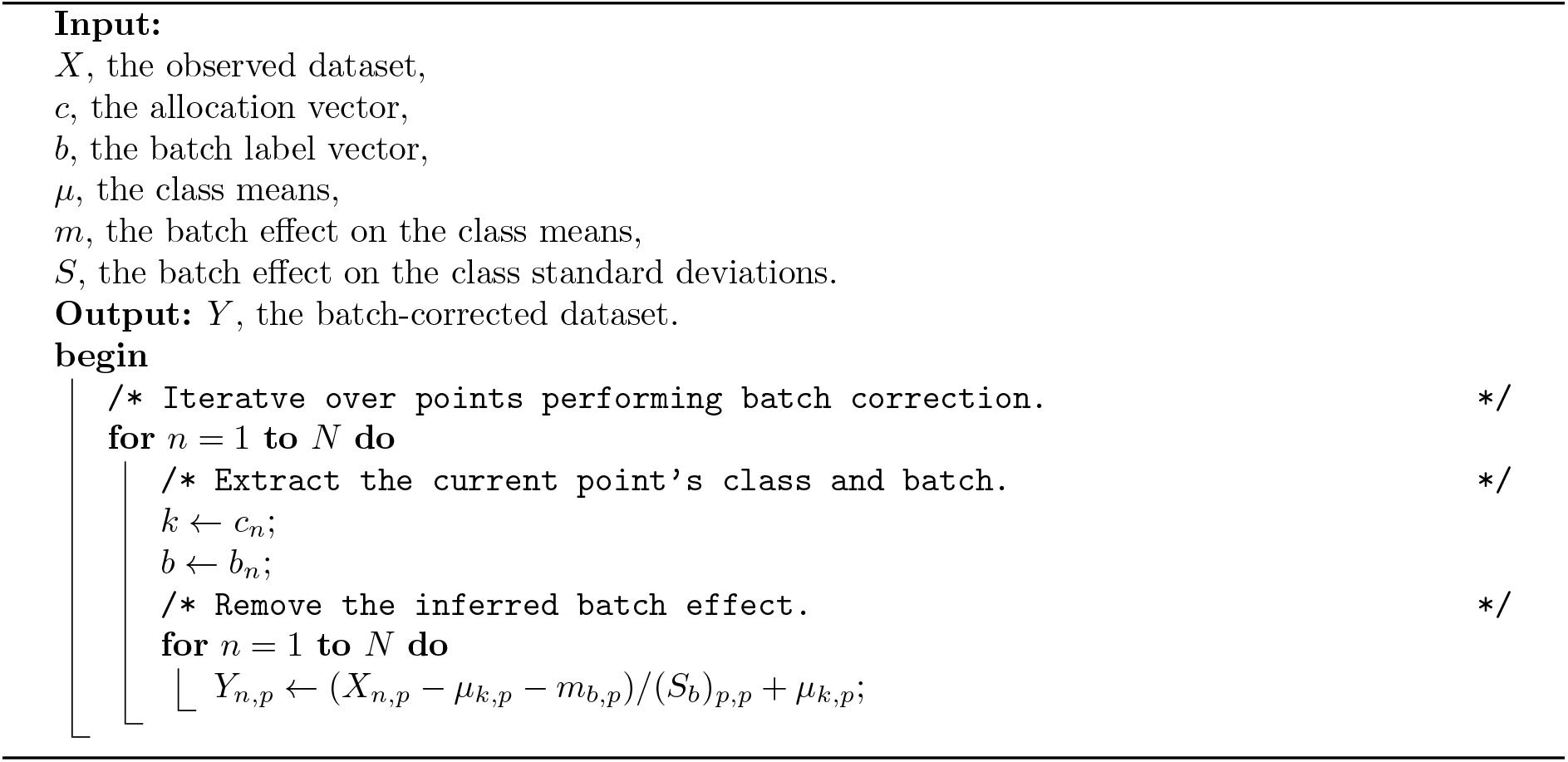

##### Algorithm 6: metropolisHastings(θ, 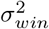, q(·))

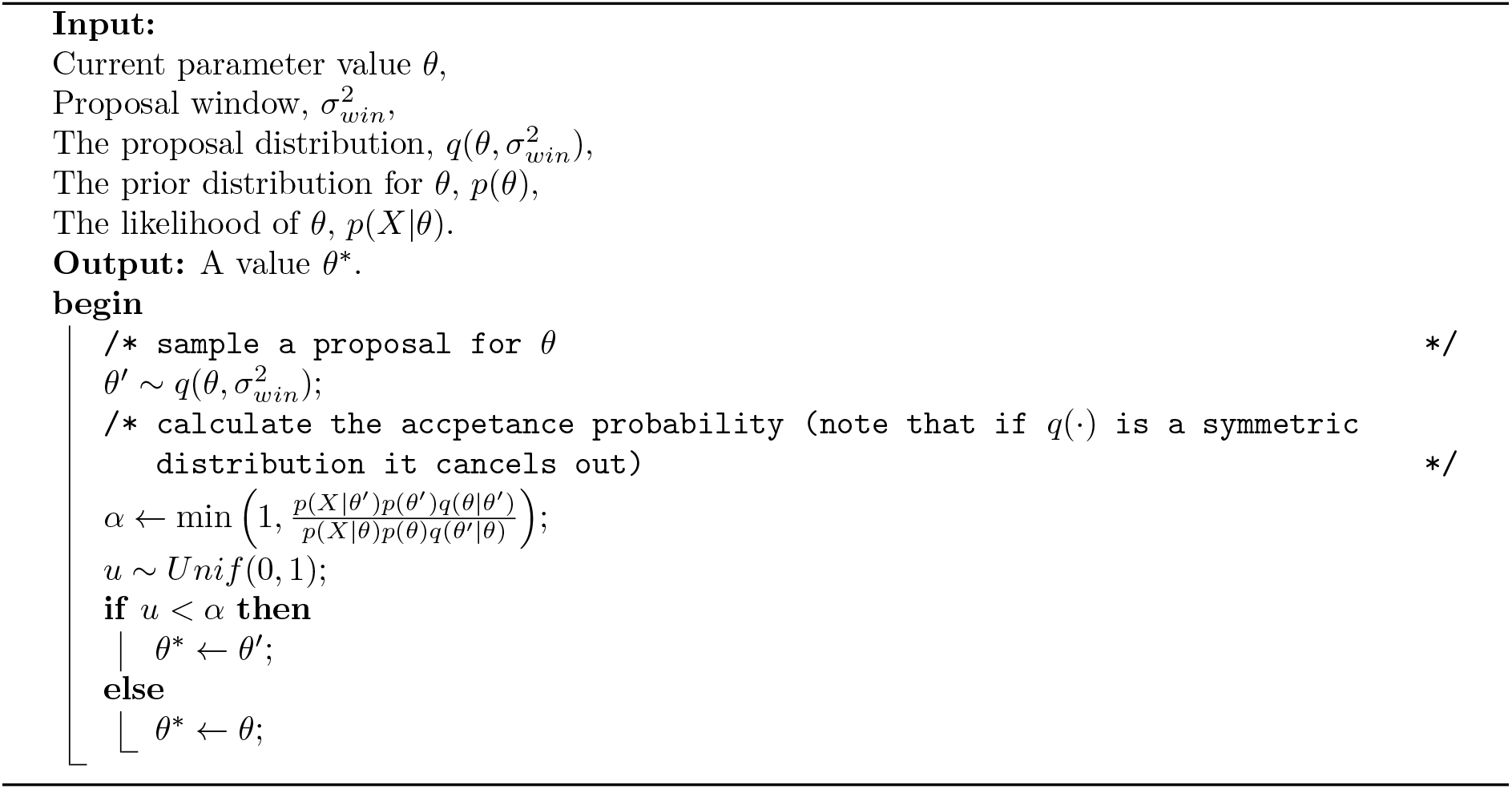

This density is symmetric and the relationship between the acceptance rate and the choice of the proposal window (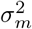 and 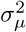) is relatively intuitive, the acceptance rate will decrease as the window increases.

The batch standard deviations have a support of (*S_loc_*, ∞). To ensure that proposed values remain in this range we use a Gamma proposal distribution with a shape of the current value divided by the rate, the rate set to some constant and a location of *S_loc_*.

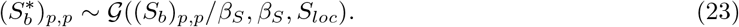

This proposal has an expected value of (*S_b_*)_*p,p*_. However, it is asymmetric and the acceptance rate increases as *β_S_* increases. We propose all *P* members of *S_b_* in each sampling step.

The class covariance matrices are the most difficult to sample. There are *P*^2^ values to propose and must be positive semi-definite. We use a Wishart proposal to satisfy this

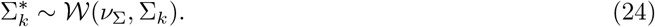

All of the proposal windows, 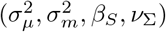, are tuned aimming to achieve acceptance rates in the range [0.1, 0.5] (Roberts and Rosenthal, 2001); if this is not possible we prioritise keeping acceptance rates above 0.1. This can involve multiple tuning runs of the sampler on each dataset.

### 4 Simulation study

We use a simulation study to test the model behaviour in examples where the generating model and the true labelling are known. We aim to explore

- the batch effects inferred by the model when none are present.
- the sampled distributions of the degree of freedom parameters in the mixture of multivariate t distributions.
- how the model behaves when there is some sort of inequality in the batches, e.g.,

– different batch sizes,
– different class representation in each batch, and
– large difference in the magnitude of batch effects.
- how the model handles misspecification.

#### 4.1 Design

Our study uses six different scenarios to test and benchmark behaviour. We use a *Base case* as the default scenario that each other scenario is a variation of. For example, the *No batch effects* scenario is the Base case with the batch means set to 0 and the batch standard deviations set to 1.0. We define each scenario by a set of parameters

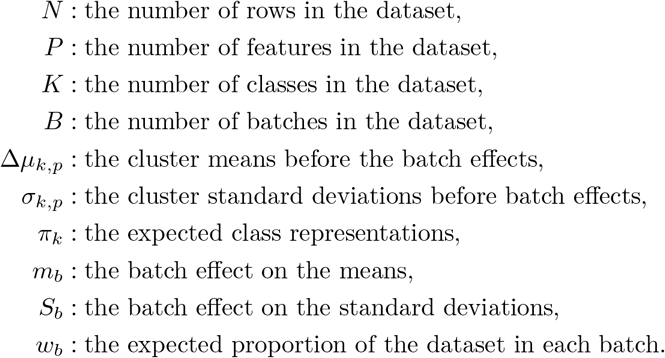

**Figure 2:**
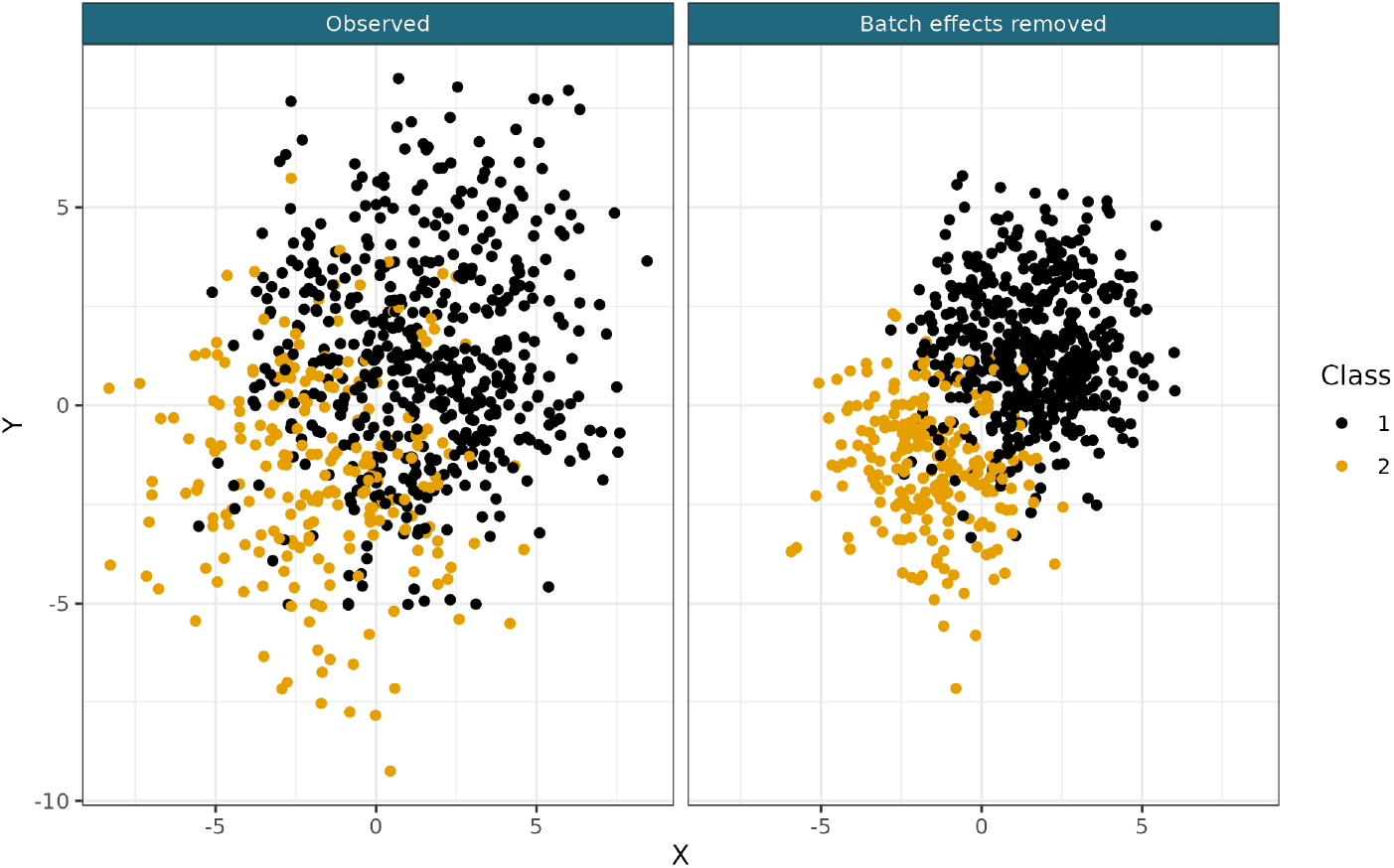
Example of a generated dataset from the Base case scenario.

We use the distance between cluster means in a single dimension, as this is the quantity of interest rather than specific values of *μ_k_*.

To generate the datasets, we first sample batch and class labels based on *w_b_* and *π_k_* respectively. The measurements for each point are then generated from a Gaussian distribution defined by these labels (except in the *multivariate t generated* scenario where the generating distribution is the eponymous distribution). We use a diagonal covariance matrix for simplicity. Each column generated randomly permutes the parameters associated with each class and batch; this means that the different columns can contain different information.

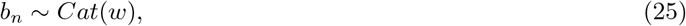

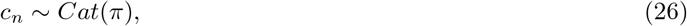

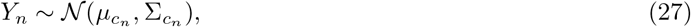

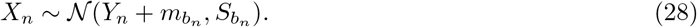

##### 4.1.1 Base case

The parameters defining each simulation in the scenario are

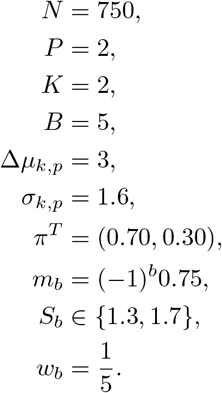

All the scenarios used these same parameters unless explicitly stated otherwise.

**Figure 3:**
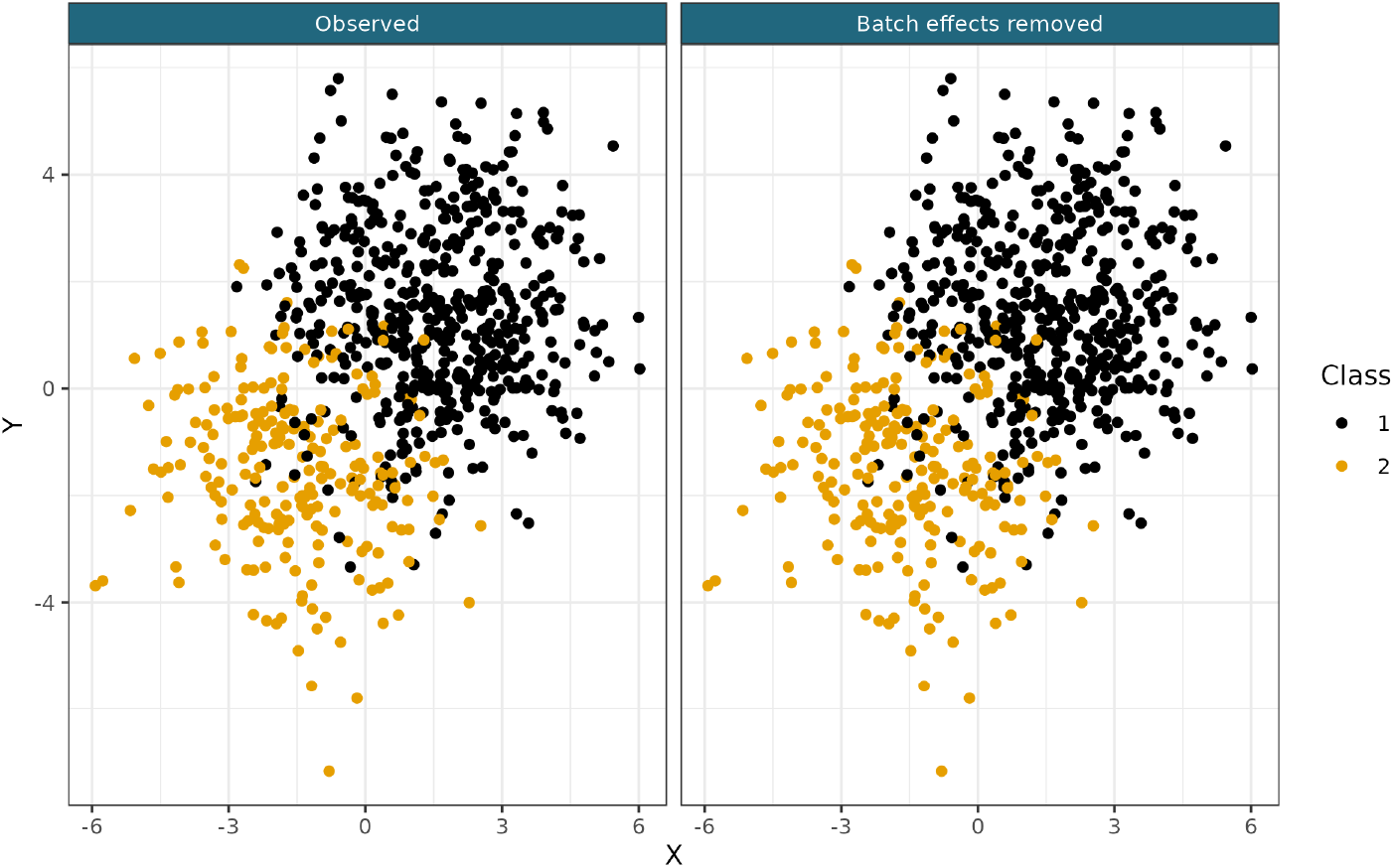
Example of a generated dataset from the No batch effects scenario. Note that the dataset is identical before and after batch-correction.

##### 4.1.2 No batch effects

This scenario is aimed at measuring the bias of the inferred batch effects. We remove the batch effects from the generating model by using values

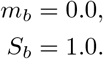

Note the inferred values of *S* are restricted to the open interval (1, ∞) in our sampler. Because of this we hope that the sampled batch scaling effect has a similar distribution across all batches rather than sampling a distribution centred on 1.0.

##### 4.1.3 Varying batch size

This scenario investigates the behaviour of the model when the batch sizes are very different.

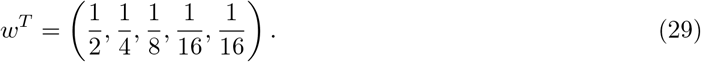

##### 4.1.4 Varying batch effects

This scenario tests how successfully the model infers to differing batch effects in each batch, different magnitudes of batch effects (with some in the tails of the prior distribution) and the direction of the batch mean shift.

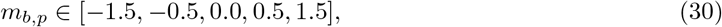

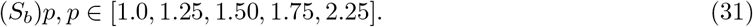

##### 4.1.5 Varying class representation across batches

In this scenario we investigate how the model responds to different expected representation of classes in each batch. This scenario might apply if the batches are collected across time and the proportion of each class in the population is expected to fluctuate. In this case the expected class proporitons vary across batches are therefore a *K* × *B* matrix,

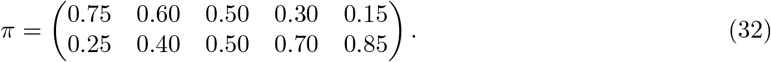

**Figure 4:**
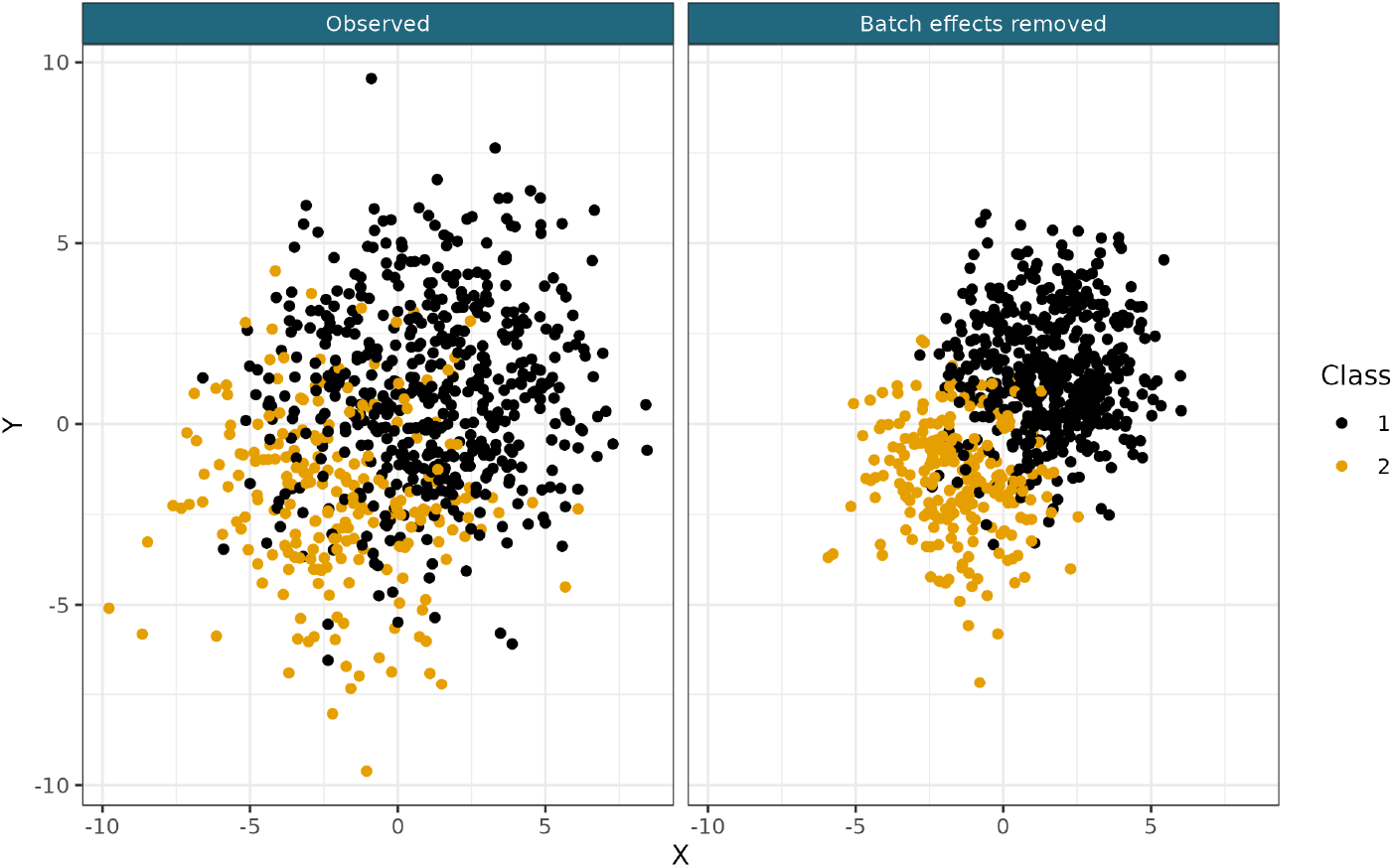
Example of a generated dataset from the Varying batch size scenario.

**Figure 5:**
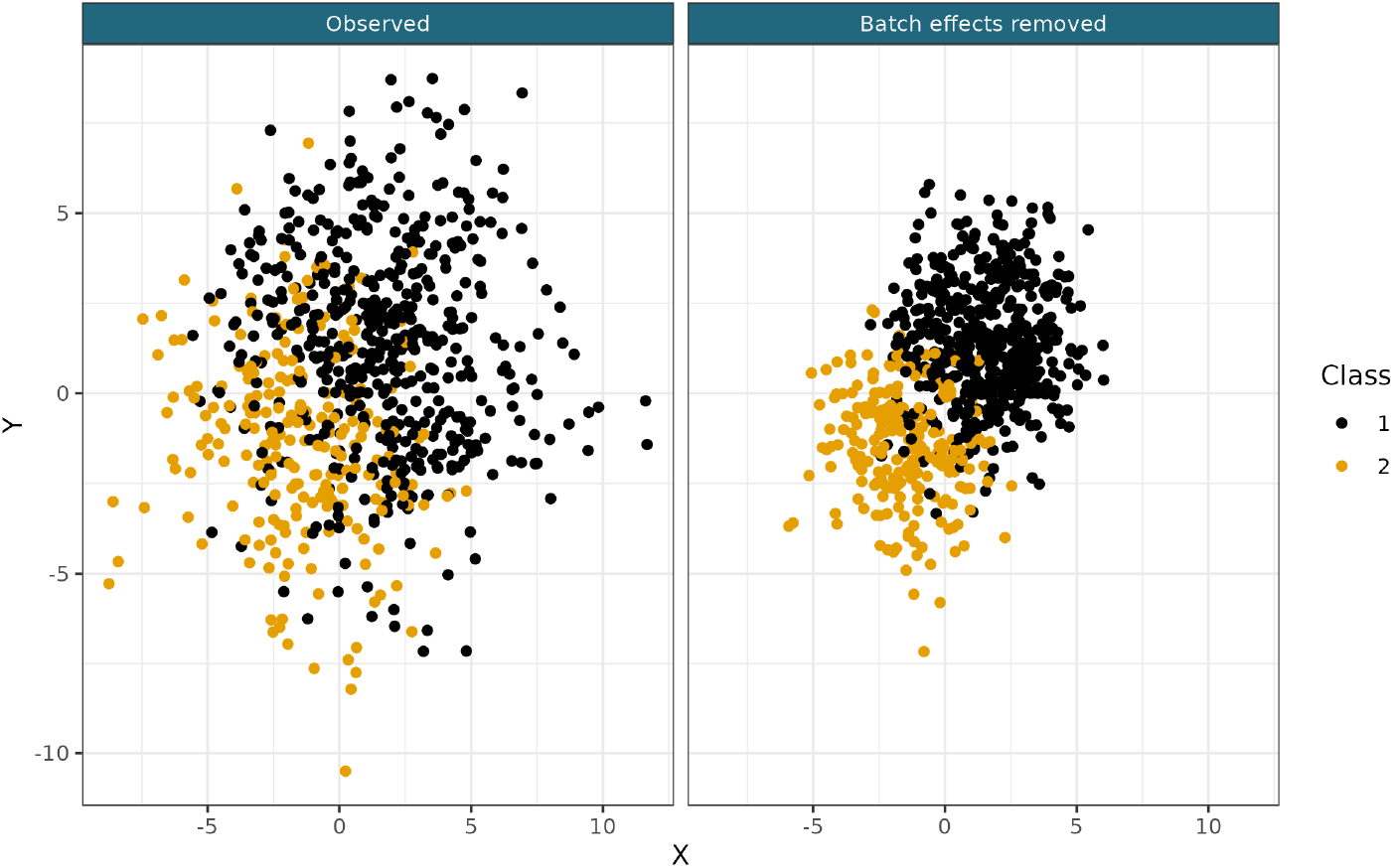
Example of a generated dataset from the Varying batch effects scenario.

**Figure 6:**
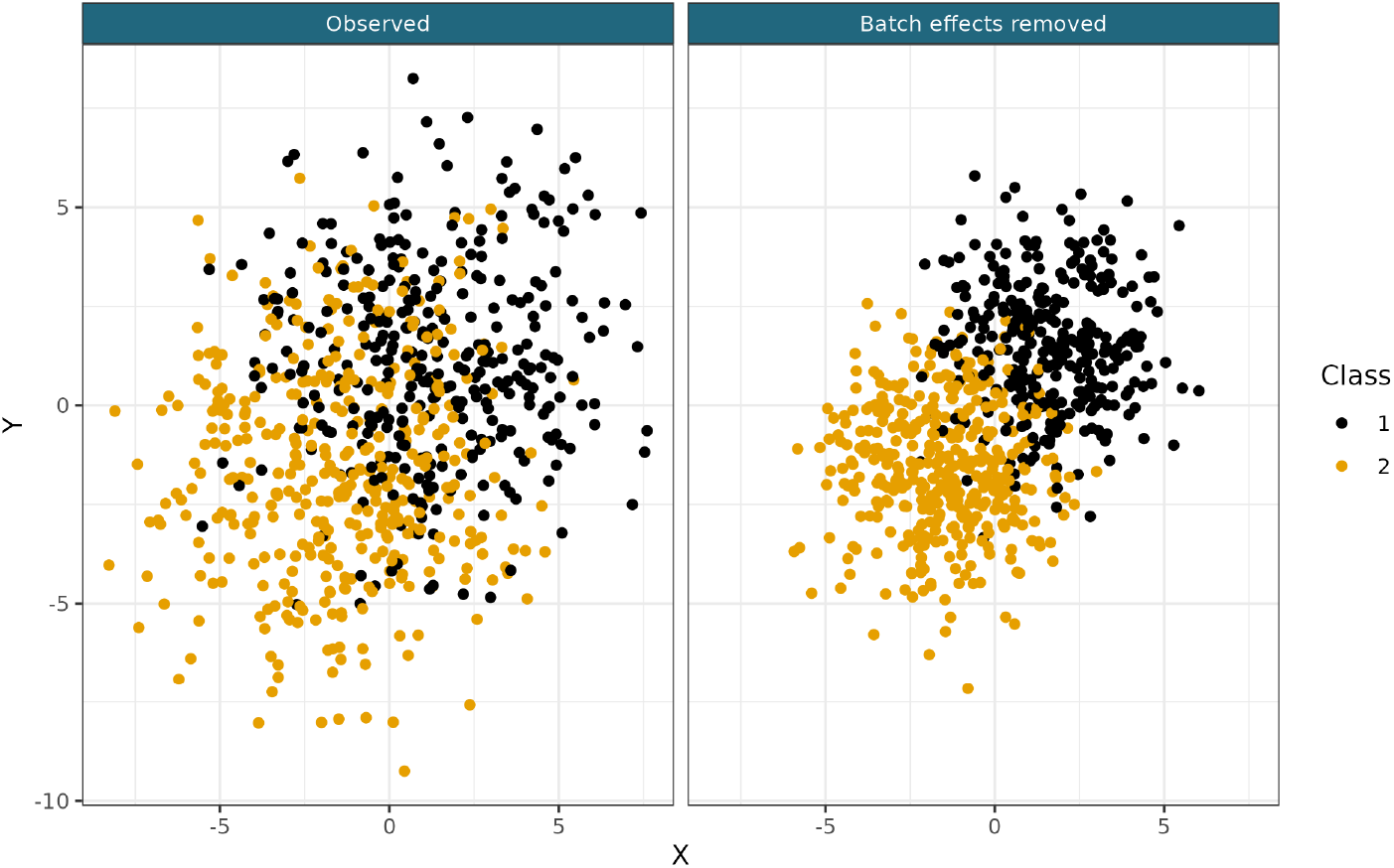
Example of a generated dataset from the Varying class representation across batches scenario.

In each batch one column of this matrix is used to sample the class membership. This introduces a dependency for *c_n_* on *b_n_*, i.e.,

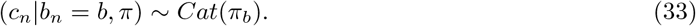

##### 4.1.6 Multivariate t generated

This scenario generates the data for each class from a multivariate t (**MVT**) distribution. This type of data is believed to be common in biology and we wish to investigate how well the model learns the degrees of freedom parameter and to compare the performance of the mixture of Gaussians model to the mixture of MVTs model.

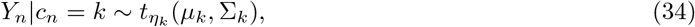

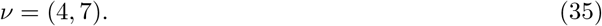

##### 4.1.7 Log-poisson generated

This scenario generates the data for each class from a Poisson distribution distribution, which is then log-transformed and has some Gaussian noise added. This type of data is believed to be common in biology where count data is so prolific and we wish to investigate how well the model behaves when it is strongly misspecified.

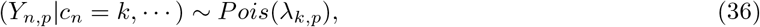

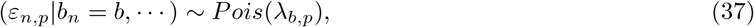

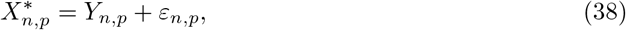

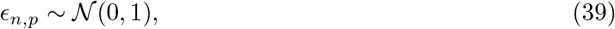

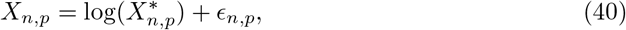

for all *n* = 1, …, *N, p* = 1, …, *P*, with *X* being the modelled data.

##### 4.1.8 High-dimensional

The data are generated from a mixture of MVN distributions as in the Base case, but more features (*P* = 15) are generated. To compensate for the additional information these contain, the distance between the class means in each feature is reduced.

**Figure 7:**
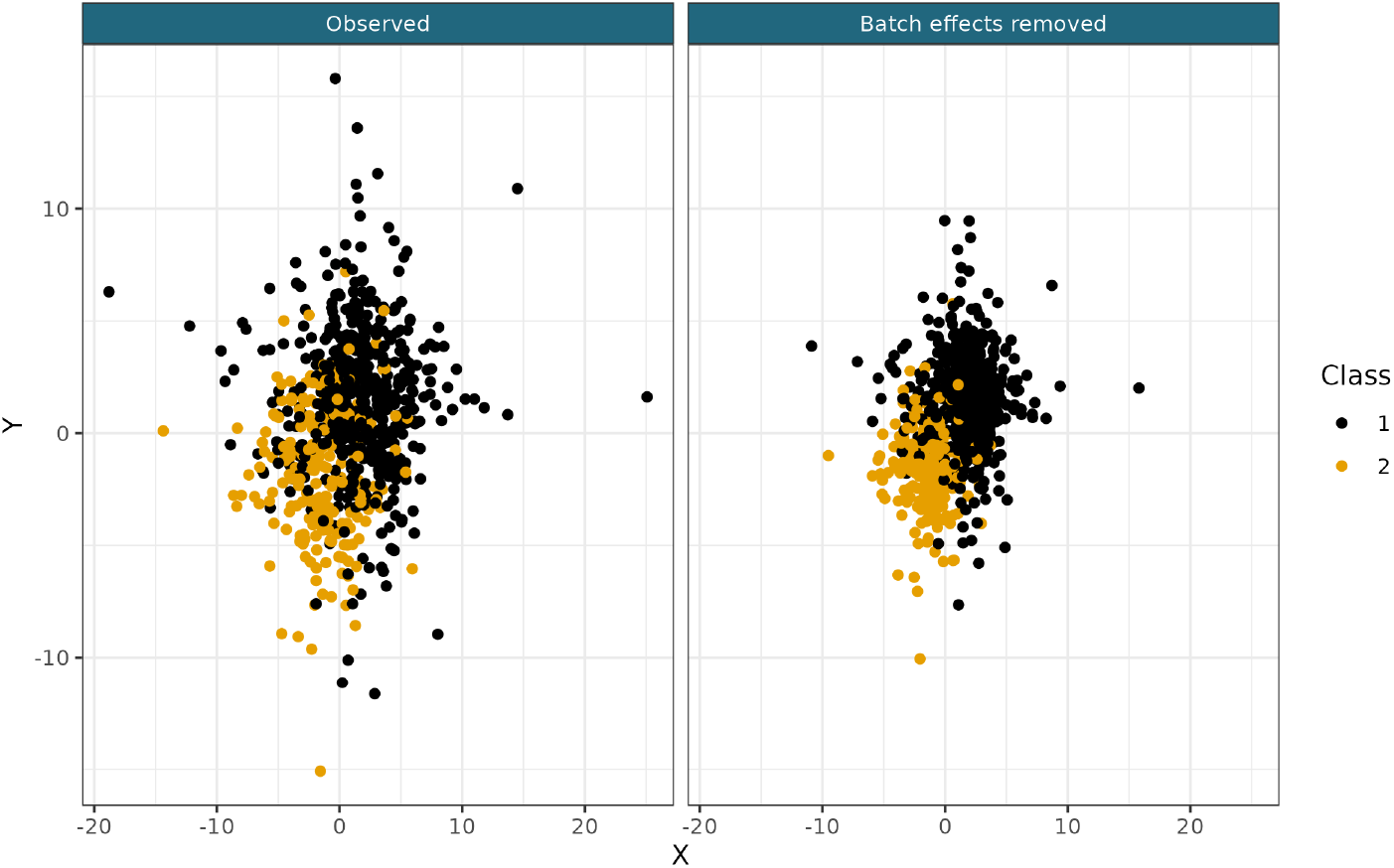
Example of a generated dataset from the MVT scenario.

**Figure 8:**
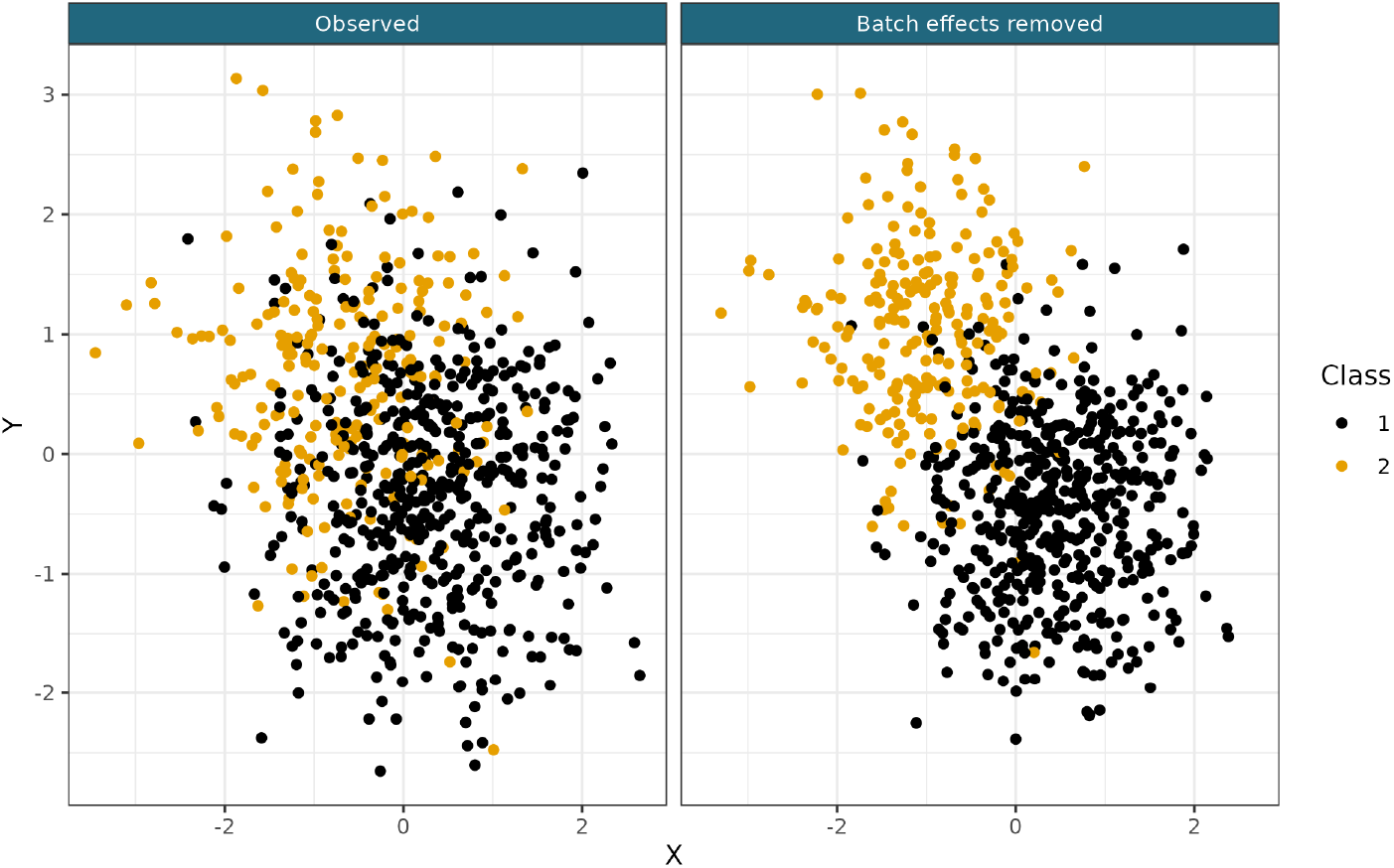
Example of a generated dataset from the Log-Poisson scenario.

**Figure 9:**
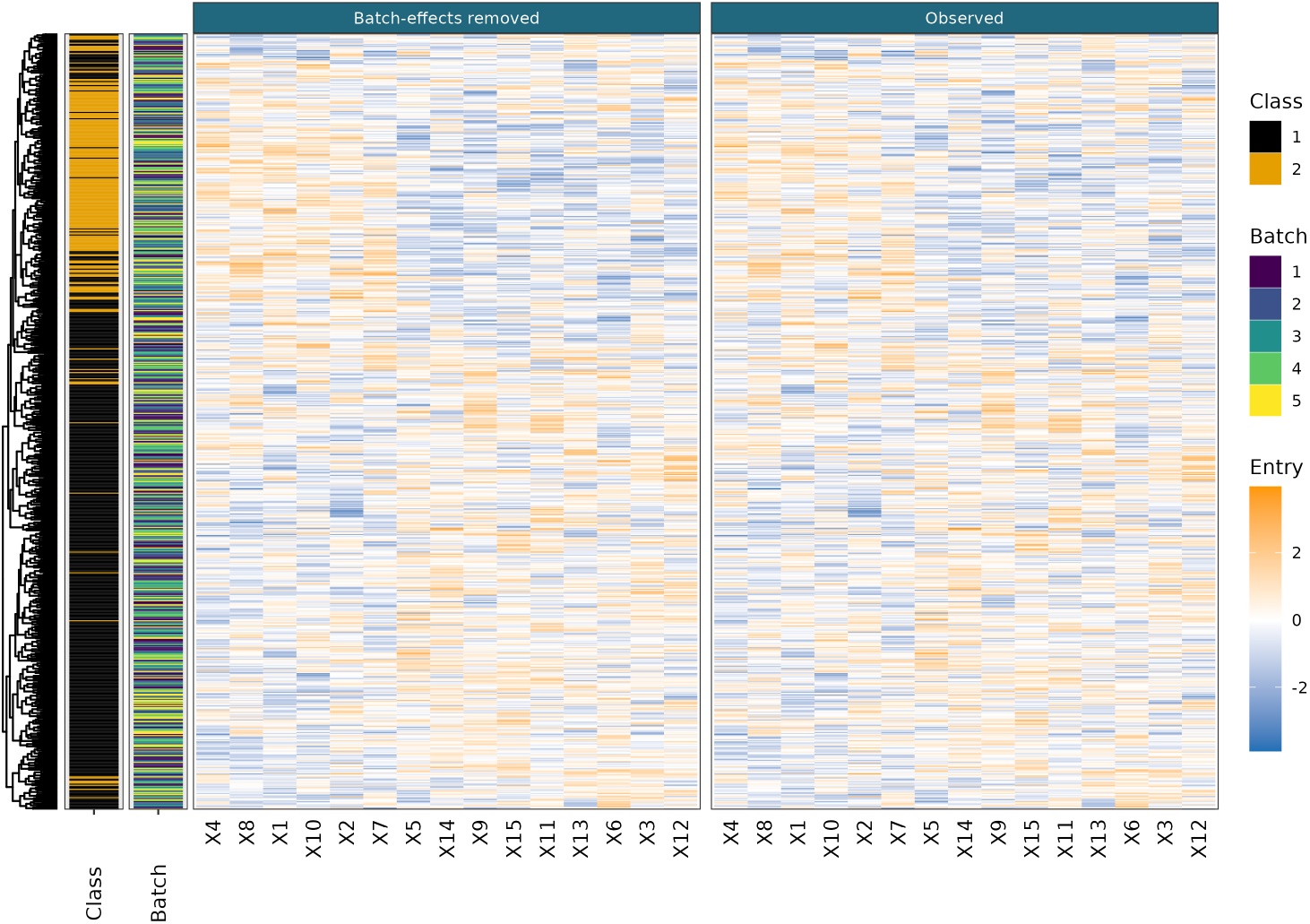
Example of a generated dataset from the High-dimensional scenario. Samples being clustered in rows, measurements in columns.

##### 4.1.9 Hardest

This scenario uses the class / batch dependency of Varying class representation across batches scenario, the data generating mechanism of the Log-poisson generated scenario and the dimensionality of the High-dimensional scenario.

#### 4.2 Additional results

### 5 Gene expression data

The batch labels, *b* = [*b*_1_*, …, b_N_*]^⊤^, *N* = 242, are generated according to one of two scenarios described below, and then used to add batch-effects to the gene expression data.

#### 5.1 Scenario a) No dependency between class and batch

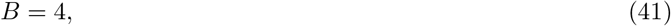

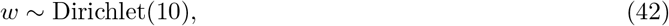

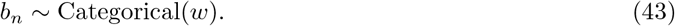

#### 5.2 Scenario b) Class and batch are dependent

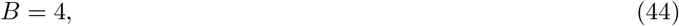

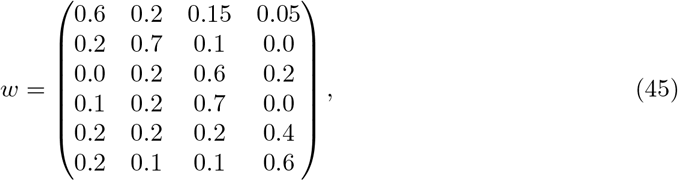

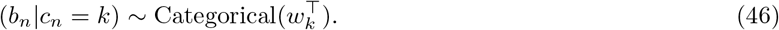

**Figure 10:**
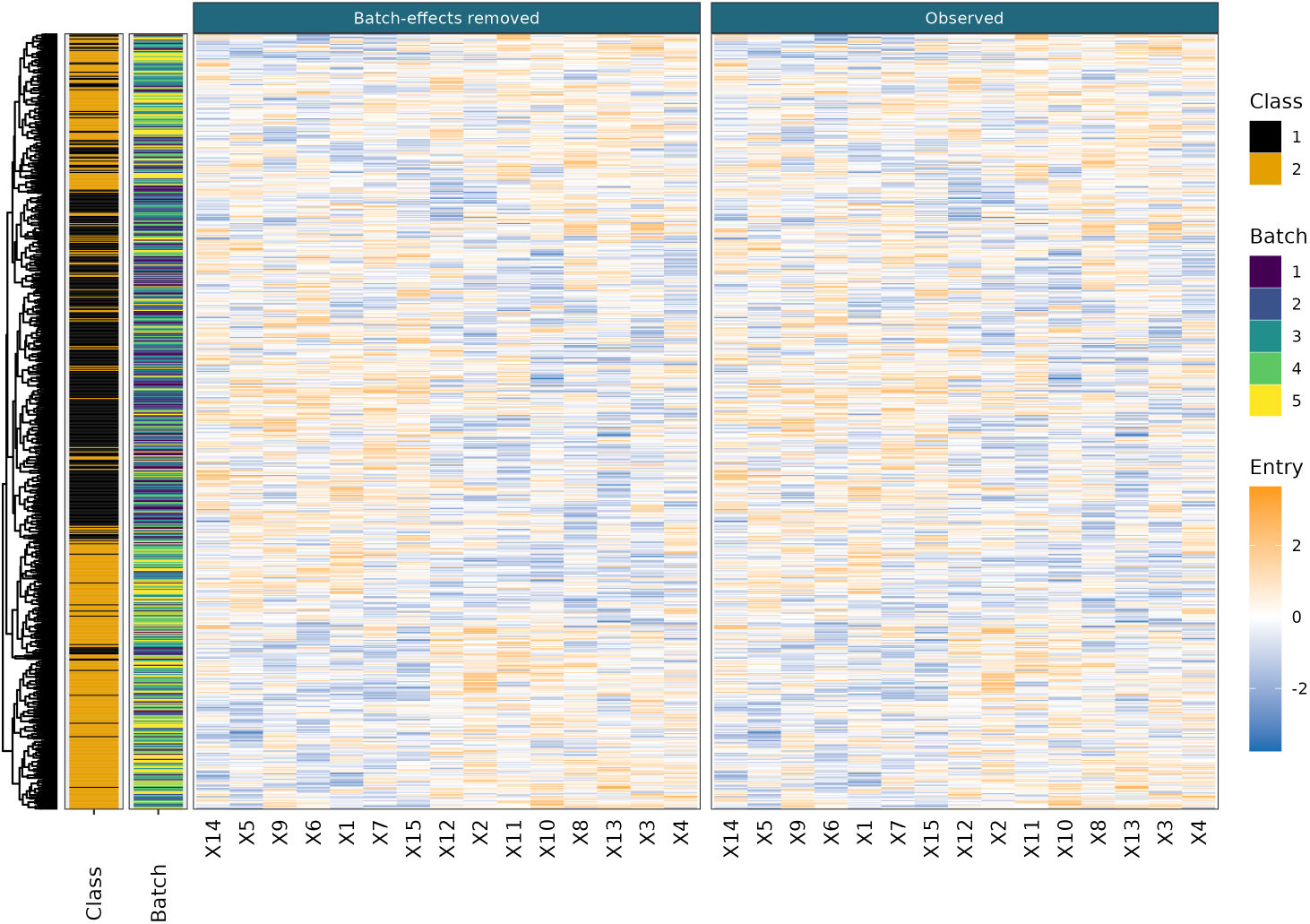
Example of a generated dataset from the Hardest scenario. Samples being clustered in rows, measurements in columns.

#### 5.3 Batch-effects

Given the batch of origin, we then simulate batch parameters

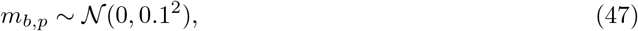

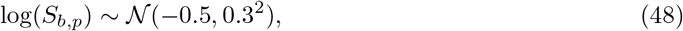

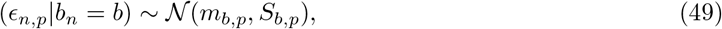

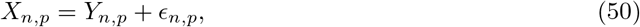

where *X* = (*X*_1_, …, *X_N_*) is the modelled data, *Y* = (*Y*_1_, …, *Y_N_*) is the gene expression data as represented in its first four principal components and *n* = 1, …, 242 and *p* = 1, …, 4.

### 6 Model convergence

For the simulated data we use the Geweke diagnostic for the complete log-likelihood after burn-in to assess within-chain convergence. We obtain a *p*-value by transforming the absolute value of the *Z*-scores with the Gaussian cumulative distribution function. We then discard all chains which have *p*-values below a threshold of 0.05. From the remaining chains we use that which has the highest median complete log-likelihood.

For the real data we visually inspect the complete log-likelihood trace plots and manually select which chains have converged to the same mode in the posterior distribution (possibly the global mode). Performing the entire process manually is feasible for the real datasets as there are less chains. An example of this process is shown in figure 12.

### 7 Dopico *et al*

Table 1 shows the seroprevalence estimate for the different methods in the data from Castro Dopico et al. (2021).

**Figure 11:**
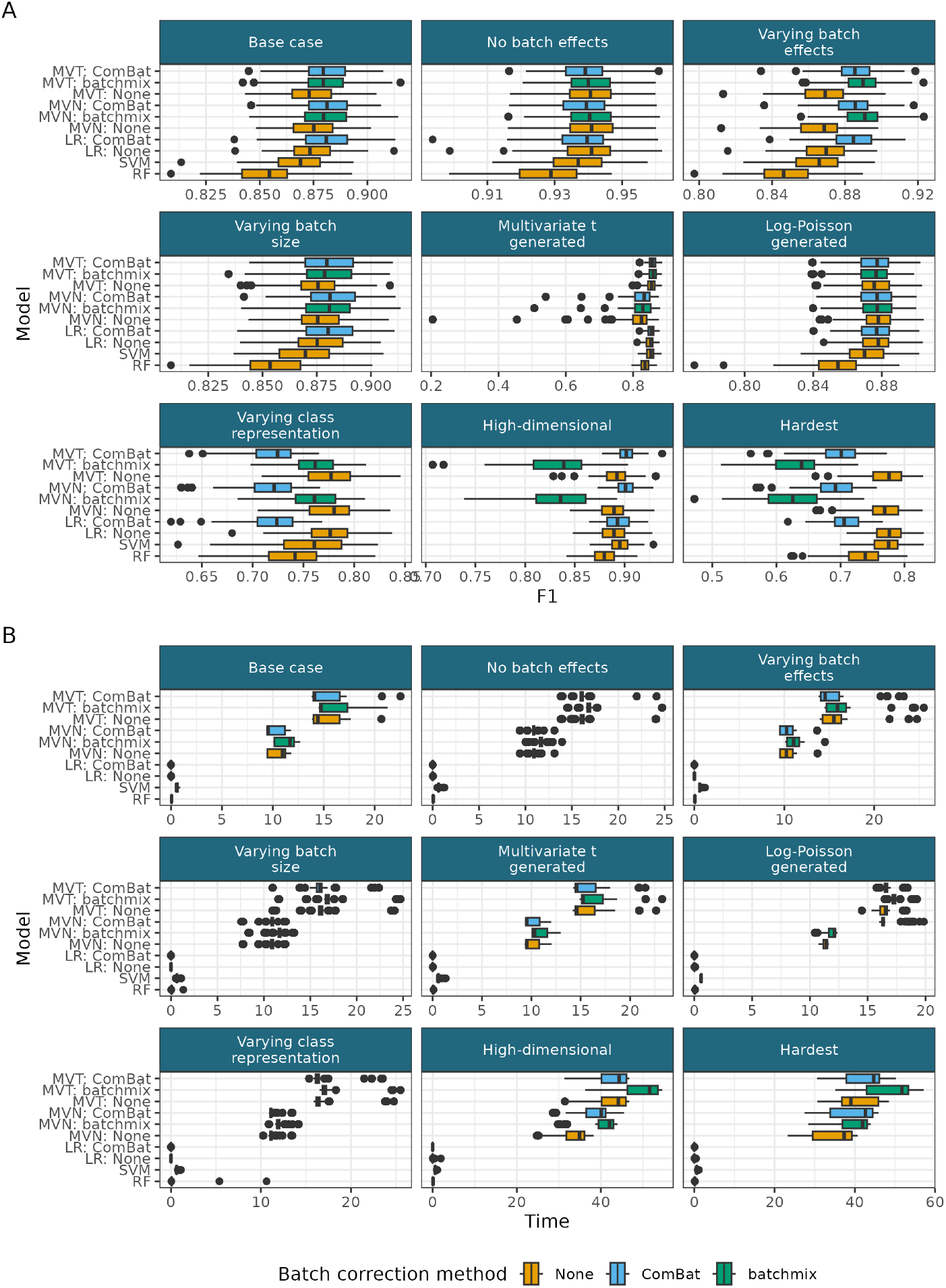
Additional results for the simulation study. A) F1 score for the unlabelled data. B) Time taken for each method.

**Figure 12:**
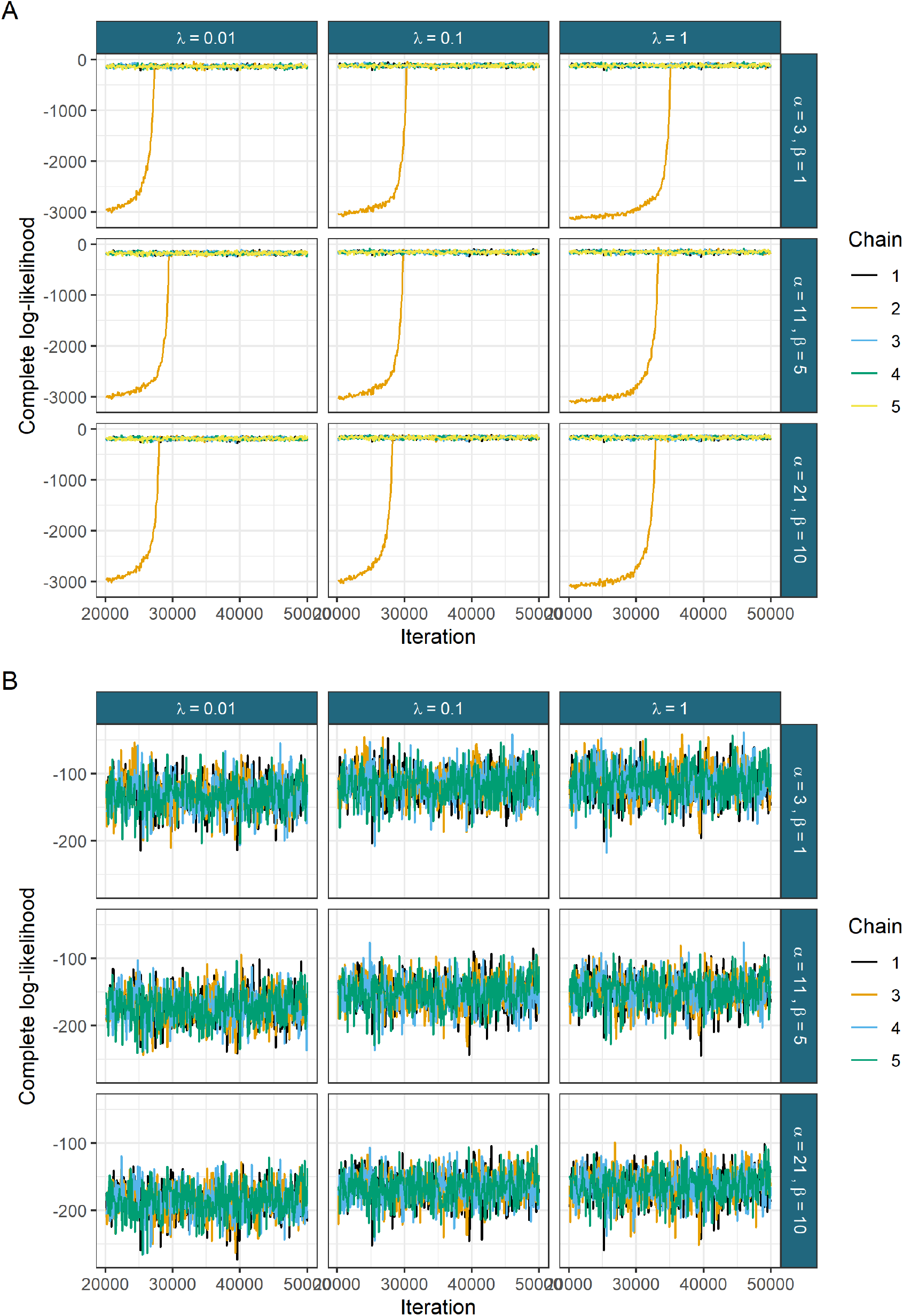
The complete log-likelihood for the MVT model in Stockholm ELISA data for A) all chains and B) the converged chains.

**Table 1:** Seroprevalence estimates across time for each method in the data from Castro Dopico et al. (2021). The highest estimates at each data are coloured orange, the lowest are coloured blue. * from Castro Dopico et al. (2021).

| Date | SVM-LDA* | Bayesian learner* | MVT | RF | SVM | LR | LR - BC |
| --- | --- | --- | --- | --- | --- | --- | --- |
| 2020/04/05 | NA | **2.35** | 1.60 | **1.00** | **1.00** | **1.00** | 2.00 |
| 2020/04/26 | **4.26** | 4.73 | 4.92 | 4.50 | **5.00** | **5.00** | **5.00** |
| 2020/05/03 | **4.33** | 5.34 | 5.74 | 4.50 | 4.50 | 5.00 | **5.50** |
| 2020/05/10 | 7.87 | **5.86** | 8.87 | 8.50 | 8.50 | 8.50 | **10.50** |
| 2020/05/17 | 7.36 | 6.28 | 8.05 | 7.50 | **4.50** | 8.00 | **8.50** |
| 2020/05/24 | 7.49 | 6.64 | 8.34 | 8.00 | **6.50** | 7.50 | **9.50** |
| 2020/05/31 | 3.54 | **6.97** | 4.15 | 4.00 | **3.50** | 4.00 | 5.00 |
| 2020/06/07 | 8.41 | **7.31** | 9.83 | 9.50 | 8.00 | 10.00 | **10.50** |
| 2020/06/14 | **6.90** | 7.75 | 7.55 | 7.00 | 7.00 | 7.00 | **8.50** |
| 2020/06/21 | **6.44** | **8.30** | 7.80 | 7.00 | 6.50 | 7.50 | 8.00 |
| 2020/07/26 | 13.20 | **11.34** | **16.72** | 15.00 | 11.50 | 16.50 | 16.00 |
| 2020/08/02 | 9.16 | **11.79** | 10.48 | 9.00 | **8.50** | 10.00 | 10.00 |
| 2020/08/09 | 11.30 | 12.15 | 13.43 | 12.00 | **8.00** | 13.00 | **13.50** |
| 2020/08/16 | 10.84 | **12.43** | 12.15 | 12.00 | **10.50** | 11.50 | 11.00 |
| 2020/08/23 | 12.97 | 12.65 | **15.55** | 15.50 | **11.00** | 15.00 | 15.00 |
| 2020/11/08 | 11.79 | **14.28** | 13.72 | 12.00 | **11.50** | 12.50 | 14.00 |
| 2020/11/15 | **14.20** | 14.47 | **18.85** | 15.50 | 14.50 | 17.00 | 18.50 |
| 2020/11/22 | **13.37** | 14.72 | **16.44** | 15.50 | 15.50 | 15.50 | 16.00 |
| 2020/11/29 | **13.20** | 14.98 | **17.36** | 15.00 | 15.00 | 16.00 | 16.50 |
| 2020/12/06 | 11.52 | **15.29** | 14.47 | 12.50 | **11.00** | 13.00 | 13.50 |
| 2020/12/13 | 15.73 | **15.64** | **19.65** | 18.00 | 16.00 | 18.00 | 19.00 |

### 8 Pseudo-ELISA data

We use the mean posterior values from a converged chain from the MVT mixture model as the parameters to generate the ELISA-like data. For the class parameters, these are:

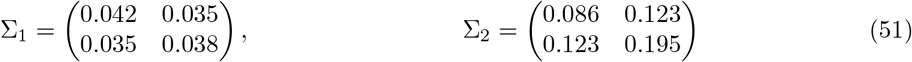

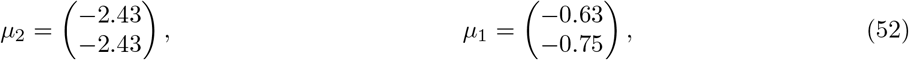

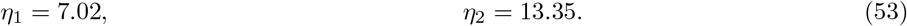

and for the batch parameters,

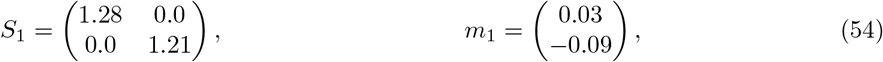

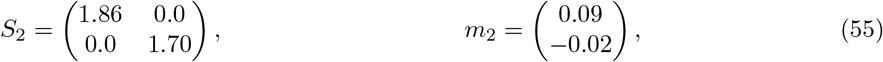

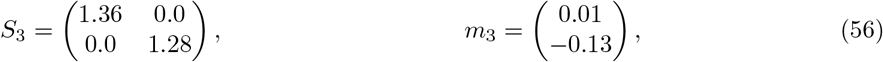

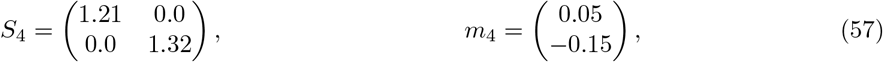

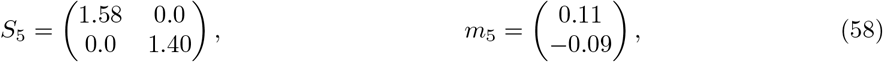

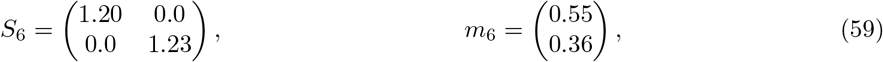

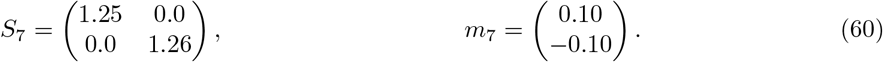

We use the predicted proportion of each batch as our batch-specific class weights,

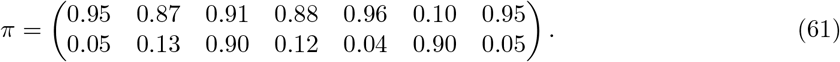

Each column corresponds to a batch and each row is the class weight. We denote the class weights within a batch (i.e., one of these columns) by π_*b*_. The probability of being drawn from a given batch is simply the observed proportion of items in each batch.

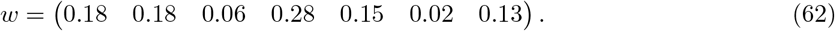

We then generate a batch and class label for each item and then observed measurements conditioning on these labels, specifically for a given item index *n*:

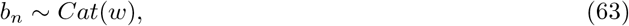

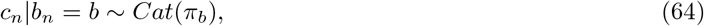

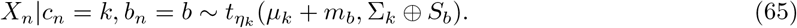

For the seronegative class (we use the label of *c_n_* = 1 for this class), the *ϕ_n_* parameter indicating if the *n^th^* item has an observed label is a Bernoulli random variable. For the seropositive class we introduce a bias to match the reality that it is more extreme observations that tend to have an observed label. To do this we find the most extreme value in each measurement, denoted *X_max_*, (note that *X_max_* is unlikely to be an observed value) and calculate the Euclidean distance between this and our observed values. We then sample *ϕ* according to:

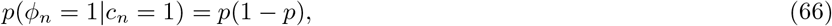

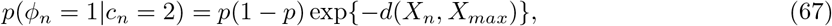

where 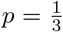. This values is chosen as the proportion of observed labels to the predicted labels is 0.332 for the seronegative class and 0.241 for the seropositive class. Our sampling process finds provides less observed seropositive labels than we have in the real data (the ratio of observed labels to true labels for the seropositive class had a mean of 0.16 across 500 simulated datasets), but we think representing the bias in the positive controls is more important than acquiring the exact proportion of training data.

